# Structure and substrate recognition by the Twin-arginine translocation (Tat) pathway core complex

**DOI:** 10.1101/2025.09.18.677151

**Authors:** J.C. Deme, O.J. Bryant, M.R.B. Batista, P.J. Stansfeld, B.C. Berks, S.M. Lea

## Abstract

The twin-arginine translocation (Tat) system is a mechanistically unique protein transport pathway moving folded proteins across membranes (*1–4*). It is found in all domains of life and is essential for plant photosynthesis and bacterial virulence(*2, 5–7*). A core complex composed of membrane proteins TatA, TatB, and TatC binds substrate proteins and then recruits additional TatA protomers to form the transport site(*8–10*). Here we present structures of resting and substrate-bound states of the Tat core complex together with biochemical and functional analysis. Substrate proteins associate with the core complex solely through their N-terminal signal peptides with their Tat targeting sequences making specific contacts with TatC and the peptide body clamped by TatB. The core complex contains highly tilted transmembrane helices that drive extreme local membrane thinning. Based on our results we propose a model for the early steps in Tat transport.

## Main Text

The bacterial cytoplasmic membrane contains two major protein export pathways, Sec and Tat, that operate in parallel to move proteins out of the cytoplasm(*2*). These pathways have been conserved in the thylakoid membrane of plant chloroplasts and in some mitochondria (and in the case of Sec, in the eukaryotic endoplasmic reticulum), making them one of the most widely distributed protein transport systems in biology(*2, 5, 7, 11–15*). Whilst the mechanism of Sec transport has been established in exquisite detail, the mechanistic basis of Tat transport remains poorly understood. The Tat pathway plays essential roles in the biogenesis of respiratory and photosynthetic machinery, formation of the bacterial cell envelope, and bacterial pathogenesis(*2, 5–7*). Substrate proteins are targeted to the Tat system by N-terminal signal peptides containing a consensus amino acid motif that includes a pair of invariant adjacent (‘twin’) arginine residues(*16–18*). Previous biochemical studies have implicated the consensus motif and residues in TatC as principal determinants of binding but the molecular nature of the interaction remains to be determined(*19–26*). In contrast to other protein transport systems that reside in ion-impermeable membranes (including the Sec apparatus), the Tat pathway transports its substrate proteins in a folded rather than unfolded state (Fig. 1a)(*7, 16, 27*). Consequently, the Tat system faces the challenge of maintaining the membrane permeability barrier while transporting substrates with wide variations in size, shape and surface properties.

**Fig. 1:**
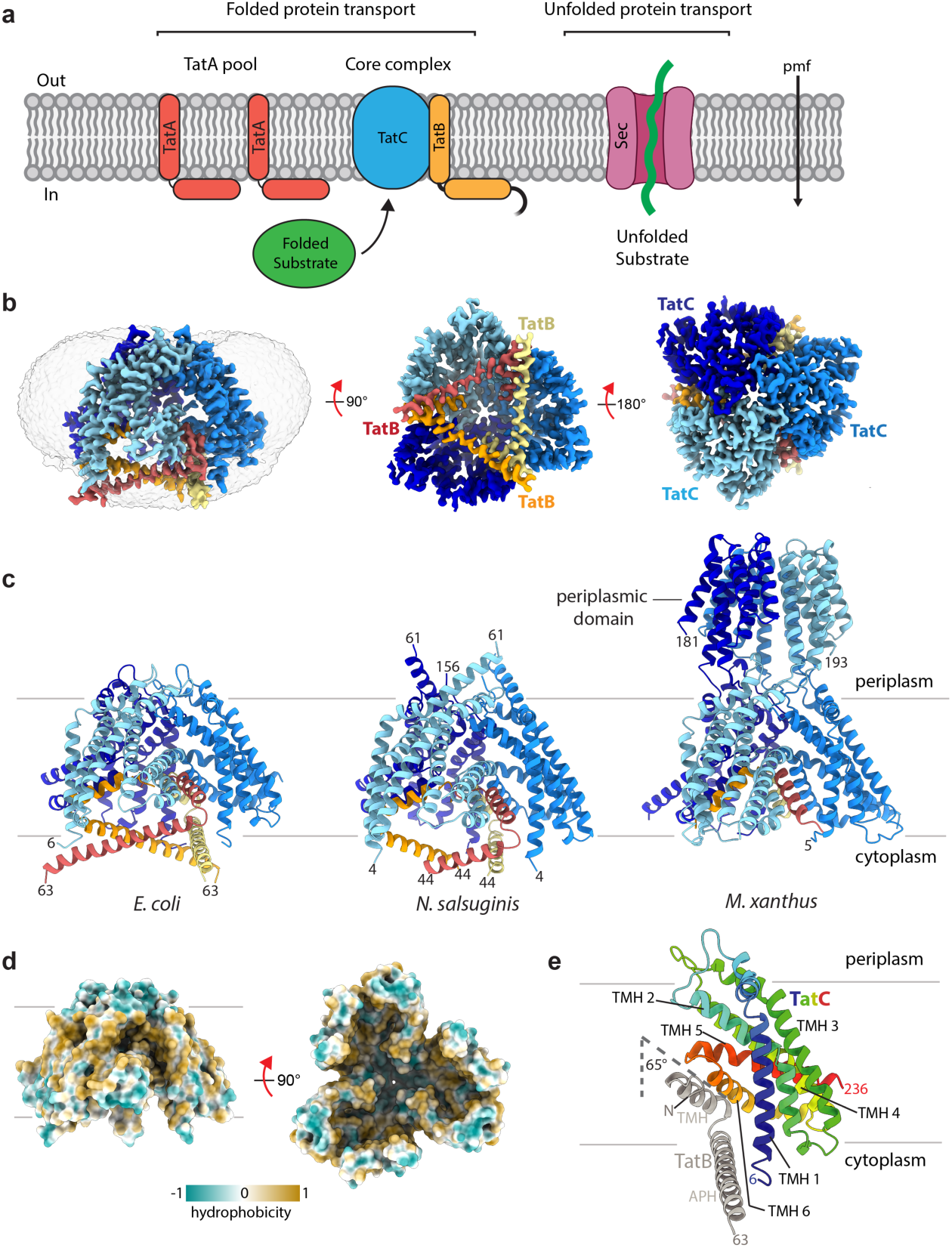
Architecture of the TatB_3_C_3_ complex. **a**, The twin-arginine translocation (Tat) system transports folded substrates across membranes and consists of a core complex minimally composed of several copies of TatB and TatC and a large pool of TatA monomers. Substrate docking to the core complex promotes pmf-dependent recruitment of multiple TatA protomers to form the active translocation site. By contrast, the Sec system found in the same membrane (right) transports proteins in the unfolded state. **b**, Cryo-EM volume of the *E. coli* TatBC complex in GDN at 3.1 Å resolution (contour level 0.175), viewed from the membrane plane (left), from the cytoplasm (middle) and periplasm (right). The detergent micelle (left; grey; contour level 0.08) indicates the position of the membrane. The APHs of the three copies of TatB form the triangle of helices visible on the base of the complex in the centre panel. **c**, Cartoon models of TatBC complexes from *E. coli* (left), *N. salsuginis* (middle) and *M. xanthus* (right) viewed from the plane of the membrane. The large periplasmic insert of *N. salsuginis* TatC is disordered and not resolved. The membrane, assigned from the position of the detergent micelle, is depicted by thin lines in this and panels d,e. **d**, Surface model of the TatC trimer within the *E. coli* TatBC complex coloured by amino acid hydrophobicity. **e,** Cartoon representation of a single TatBC heterodimer from within the *E. coli* TatBC complex. TatC is coloured from blue at the N-terminus to red at the C-terminus, and TatB is coloured grey. The angle the TatB TMH makes with the membrane normal is shown.

In most organisms the minimal components of the Tat system are the three integral membrane proteins TatA, TatB and TatC(*28–30*) (Fig. 1a). TatA and TatB are structurally related but functionally non-identical proteins that both contain a single N-terminal transmembrane helix (TMH) followed by an amphipathic helix (APH)(*31–33*). TatC contains six transmembrane helices(*21, 34*) and together with TatB forms a multimeric core complex that engages with the signal peptides of substrate proteins(*8, 22, 23, 35*). TatA protomers may also be part of this core complex(*35, 36*) but have been thought to be peripherally associated in the resting state(*37, 38*).

Docking of a substrate protein to the core complex triggers a protonmotive force (pmf)-dependent recruitment and oligomerisation of many additional TatA subunits to form the translocation site through which the folded substrate is transported(*9, 10, 19, 39*). Tat transport is not thought to utilise a conventional protein conducting channel but instead the TatA oligomer is proposed to perturb the membrane bilayer structure to allow transmembrane movement of the substrate protein(*31, 40–43*). Completion of transport is followed by disassembly of the translocation site and removal of the signal peptide from the substrate protein by signal peptidase.

The architecture and oligomeric state of the Tat core complex and the mechanism by which the Tat machinery recognise substrates are not understood. To address these questions, here we here use cryo-EM to determine structures of the Tat core complex, both in isolation and with bound substrate proteins.

## The TatBC complex

We initially focused on Tat core complexes containing just TatB and TatC, to avoid possible heterogeneity associated with the presence of TatA, and targeted the prototypical TatBC complex from *Escherichia coli* for structure determination by single particle cryo-EM. However, this proved to be challenging due to its small size and minimal features outside the solubilizing detergent micelle. We thus also targeted two atypical Tat complexes from species *Nitratifractor salsuginis* and *Myxococcus xanthus*, which contain large inserts in different TatC periplasmic loops that might aid structure determination. Additional protein density outside the detergent micelle was indeed visible in 2D classes from single-particle cryo-EM data of the TatBC complexes of both species (Extended Data Fig. 1a), and the *M. xanthus* insert folded to form a homotrimer for which we determined a structure by X-ray crystallography (Extended Data Fig. 1b).

We eventually generated 3D reconstructions and built models of the TatBC complexes of all three species at resolutions between 2.6 and 3.1 Å (Figs. 1b,c and S1-3). These models revealed a common (TatBC)_3_ organisation, in which the complex is formed from a trimer of TatBC heterodimers. The subunit stoichiometry of the TatBC complex was previously unclear and generally overestimated relative to the actual size of the complex revealed here(*3, 8, 22, 36, 44*). The periplasmic insert in TatC in the *M. xanthus* complex is ordered and folded as in the crystal structure (Figs. 1c and S1c) but is disordered in the *N. salsuginis* complex (Fig. 1c and S1a). The remaining parts of TatC could be built in their entirety except for a small number of residues at the polypeptide termini and in some inter-helix loops. In the better-ordered *E. coli* and *N. salsuginis* complexes, the TMH and APH of TatB were resolved (in the *M. xanthus* complex, only the TM could be built), suggesting that residues C-terminal to the APH are disordered or mobile relative to the membrane-anchored portion of the complexes. Unless otherwise indicated, we henceforth specifically reference the *E. coli* TatBC complex, in which TatB residues 1–63 and TatC residues 6–236 can be modelled, but the three complexes are highly similar (Fig. S1d)

The three copies of TatC within the TatBC complex each have a ‘glove’ shape and are arranged with their concave faces directed to the centre of the complex and their convex faces facing out towards the lipid bilayer. Compared to previous *A. aeolicus* TatC monomeric crystal structures(*21, 34*), the TatC proteins in the TatBC complex exhibit a subtle closure of the C-terminal end of the TatC ‘glove’, leading to an increase in concavity (Fig. S1e). The three TatC molecules pack tightly at the periplasmic side of the membrane, forming an inverted cup-like structure in which the largely hydrophobic interior surface is open to the cytoplasm (Fig. 1d). The observed TatC-TatC contacts include residues previously identified as important for core complex function or assembly(*23, 45*) (Fig. S1f,g). In the *M. xanthus* TatBC complex the TatC interactions are augmented by the contacts between the inserted periplasmic domains (Figs. 1c and S1c).

The TMH and APH of TatB in the core complex encompass the same span of residues as in the isolated TatB protein previously characterised by NMR(*32*), except for the small α3 helix identified by NMR, which now forms an extension to the APH (Fig. S4a,b). The TatB helices are rearranged in the context of the full TatBC complex, compared to the isolated TatB protein (Fig. S4b). Each TatB subunit is intercalated between a pair of TatC subunits (Figs. 1b,e,2a).

This arrangement is primarily stabilized by interactions between the transmembrane helix (TMH) of TatB and two regions of adjacent TatC molecules: TMH5 and the loop leading into TMH6 of the counterclockwise-adjacent TatC (as viewed from the cytoplasm), and TMH1 of the clockwise-adjacent TatC (Figs. 2a and S5a). This arrangement was anticipated by earlier studies that had implicated both ends of the TatC molecule in interactions with the TatB TMH(*21, 23, 35*).

**Fig. 2:**
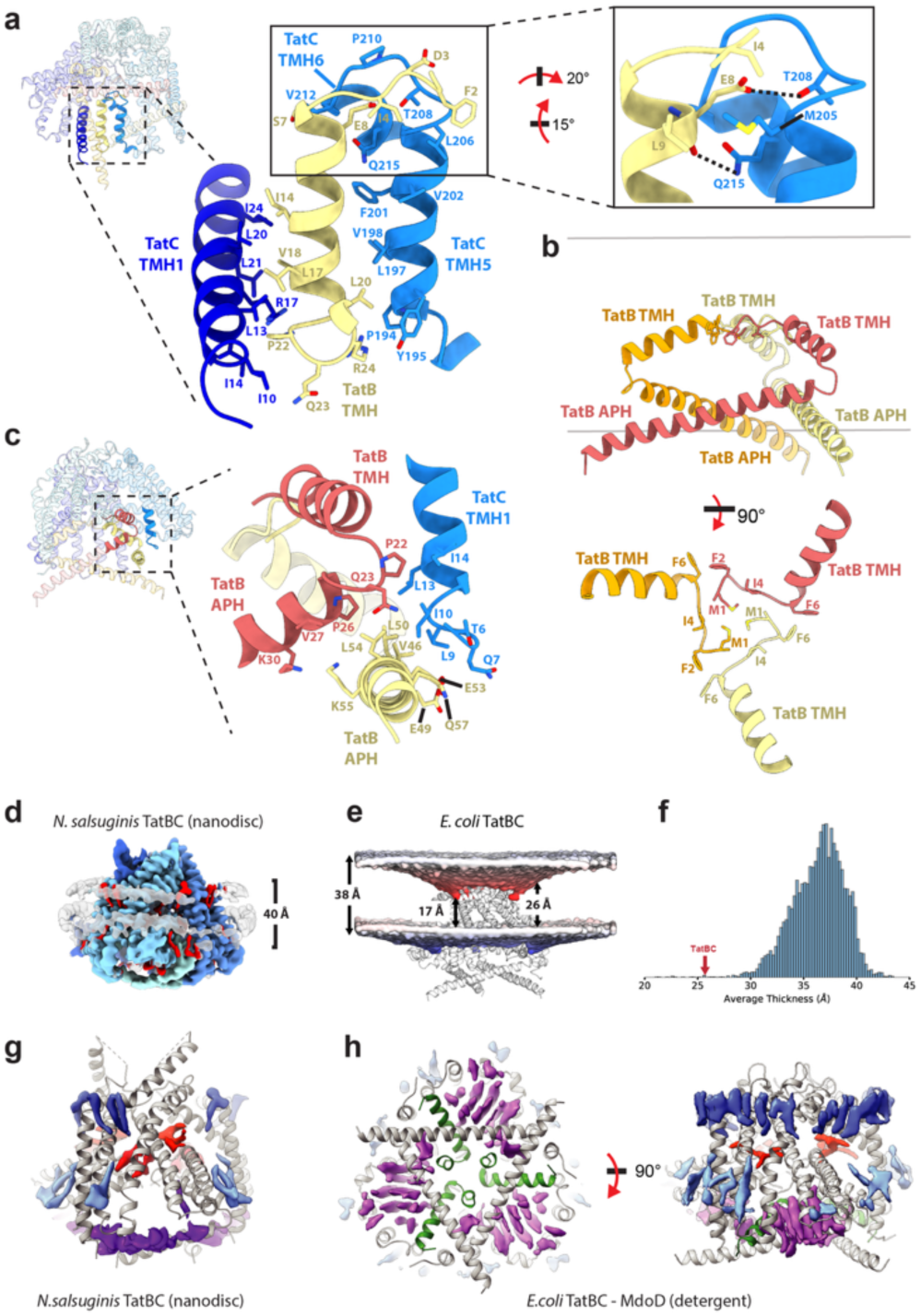
TatBC complex assembly and membrane thinning. **a**, Structure of an *E. coli* TatC-TatB-TatC interface. Shown are TMH1 of the first TatC monomer (dark blue), TMH5 of the second TatC monomer (light blue), and the TatB TMH (yellow). The zoomed in view (inset panel right) highlights the interactions found at the previously proposed ‘polar cluster site’. Hydrogen bonds are depicted as dashed lines. **b**, Structure of the TatB cage formed by the three copies of TatB (coloured in red, orange and yellow). (Top) View from the membrane plane. (Bottom) View of the three TatB TMHs viewed from the cytoplasm. **c**, Structure of an *E. coli* TatB-TatB-TatC interface. TatC TMH1 is shown in light blue, two TatB copies are shown in red and yellow. **d**, Surface representation of the mean positions of the lipid phosphates, highlighting the membrane thinning induced by the TatBC complex. **e**, Distribution of average membrane thickness values for the annular shell (within 7 Å) around the membrane proteins in the MemProtMD database. The average thickness for TatBC is highlighted. **f**, The TatBC complex thins a membrane-mimetic bilayer. Cryo-EM volume of the *N. salsuginis* TatBC complex in a lipid nanodisc at a resolution of 3.1 Å, viewed from the membrane plane (contour level 0.222). Lipid densities are coloured red and the helical nanodisc belts grey. **g**,**h**, Lipid/detergent densities visible in the cryo-EM volumes of (**g**) the *N. salsuginis* TatBC complex or (**h**) an *E. coli* TatBC–substrate complex. The densities are coloured according to their position in the complex: either vertically oriented at the base of the TatC cup (purple), or horizontally oriented higher in the TatC cup (red), or on the outside of the complex in either the periplasmic (dark blue) or cytoplasmic (light blue) leaflet of the membrane.

The TatB APHs form a triangular arrangement at the cytoplasmic face of the complex across the otherwise open TatC cup (Fig. 1b) rather than interacting with the bilayer around the complex. In this layout the N-terminal end of each APH packs against the C-terminal end of the neighbouring TatB APH and the N-terminal end of TatC TMH1 (Fig. 2b,c). Considered together, the ordered portions of the TatB proteins form a cage-like structure (Fig. 2b) contained within the TatC trimer (Fig. 1c).

The N-terminal end of the TatB TMH contains the highly conserved polar residue E8. This amino acid was previously inferred to stabilize the association of TatB with TatC through interaction with a ‘polar cluster site’ on TatC formed by M205, T208 and Q215(*35*). Our TatBC structure reveals that although the side chain of TatC T208 forms the anticipated hydrogen bond with the side chain of TatB E8, the side chain of Q215 hydrogen bonds to the main chain oxygen of E8 rather than making the predicted interaction with the side chain of this residue (Fig. 2a). TatC M205 does not directly contact TatB E8 but packs between TatB L9 and I4. The TatC molecule that provides the polar cluster interactions makes many further contacts to the E8-containing face of the TatB TMH through a track of hydrophobic residues along the face of TatC TMH5, the TMH5-6 loop, and the N-terminal end of TMH6 (Fig. 2a). The other side of the TatB TMH is stabilised by a series of hydrophobic interactions with TMH1 of the other neighbouring copy of TatC (Fig. 2a,c). These include previously identified contacts between TatC L21 and TatB V18(*35*) (Figs. 2a, S1a) and interactions with TatB made by TatC I10 and I14(*23*) (Figs. 2a,c, S1a).

The TMHs of both TatB and TatC are highly tilted relative to the membrane normal (Figs. 1e, 2b) rather than the near-vertical arrangement suggested by simulations of the TatC monomer in lipid bilayers(*21, 34*). In the case of the TatB TMHs, the N-termini of the TatB molecules pack together within the TatC cup at the midplane of the transmembrane portion of the complex (Fig. 2b) with the TMHs projecting outwards towards the cytoplasmic end of the TatC-TatC interfaces at an angle of 65° relative to the membrane normal (Fig. 1e). This extreme TMH tilting is matched by TMH 5 and 6 of TatC which also penetrate only halfway across the membrane-spanning part of the complex from the cytoplasm (Fig. 1e).

Due to their extensive TMH tilting, TatBC complexes are highly compressed along the membrane normal (Figs. 1c and S4c-e) suggesting that the surrounding membrane will be drastically thinned. This idea is supported by molecular dynamics (MD) simulations of the lipid bilayer around the *E. coli* TatBC complex that show thinning of the membrane from 38 Å (phosphate-to-phosphate distance) in the bulk phase to an average of 26 Å adjacent to the protein (Fig. 2d). This level of membrane thinning is extreme relative to other α-helical membrane proteins in MemProtMD, a database of membrane proteins in bilayers (Fig. 2e).

The MD simulations indicate that the highly compressed structure of TatBC observed in the cryoEM structures is stable in a membrane environment and thus unlikely to be an artefact arising from detergent solubilization (Fig. 2e). To confirm this experimentally we showed that the *N. salsuginis* TatBC complex has the same structure in the membrane-like environment of a lipid nanodisc as it does in a detergent micelle (RMSD 0.8 Å for all CA pairs; Figs. 2f and S6a-c). The lipid bilayer around TatBC in the nanodisc is ∼40 Å deep (complete bilayer width) compared with the ∼60 Å width of the *E. coli* cytoplasmic membrane(*46*) consistent with the hypothesis that TatBC thins the surrounding bilayer.

Both the nanodisc-reconstituted *N. salsuginis* complex (Fig. 2g) and the detergent-solubilised *E. coli* complex (Fig. 2h) reveal many ordered lipid/detergent molecules around and within the TatBC complex. Some of these densities stack against the tilted TMHs of TatB and TatC (blue densities in Fig. 2g,h). Other lipid densities are observed within the complex in the cavities created by the concave faces of the TatC molecules (purple densities in Fig. 2g,h). Finally, a pair of lipids per TatC subunit are oriented horizontally (relative to the plane of the bilayer) and packed within the periplasmic end of the TatC cup, sandwiched between TMH6 of one TatC subunit and the concave surface of the neighbouring TatC molecule (red densities in Fig. 2g,h). MD simulations of TatBC in a membrane environment demonstrated accumulation of lipid at the same locations (Fig. S6d). This indicates that the interior of the TatBC complex is occupied by lipids arranged in a markedly different manner from those in the surrounding membrane bilayer. The considerable amount of lipid within the complex and the sensitivity of the complex to detergents harsher than glyco-diosgenin (GDN, used to purify the complexes presented herein) and digitonin(*8, 47*) suggest a role for lipids in stabilising the complex.

## Structural basis for signal peptide recognition by the TatBC complex

Tat signal peptides are composed of a positively charged n-region that includes the pair of adjacent arginine residues, a hydrophobic h-region, and a polar c-region that contains a cleavage site for signal peptidase(*16*) (Fig. 3a). In bacterial Tat signal peptides, the pair of adjacent arginine residues are part of a larger S-R-R-x-F-L-K consensus motif(*16, 27*).

**Fig. 3:**
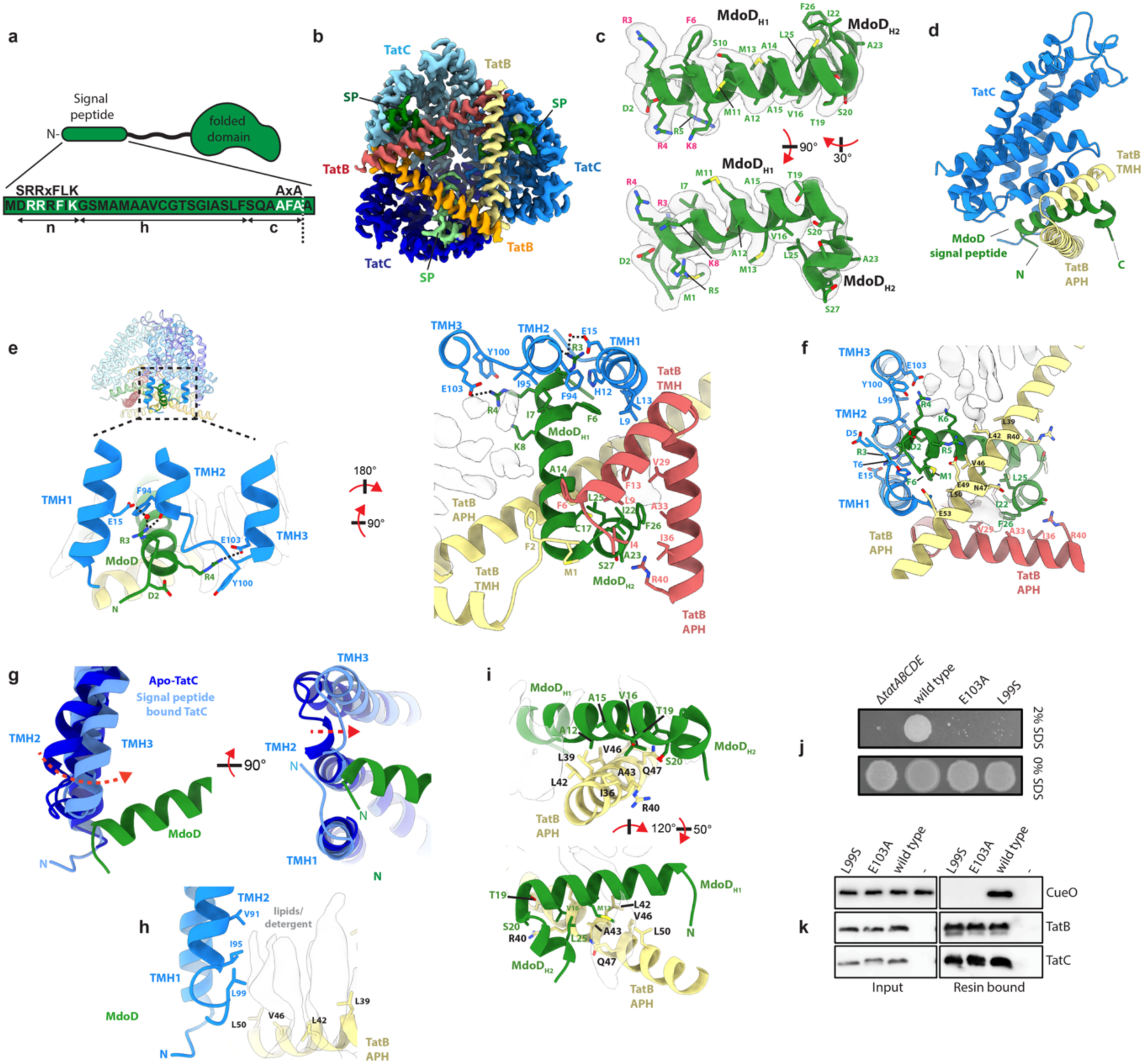
Signal peptide recognition by the TatBC complex. **a**, The features of a Tat signal peptide shown schematically (above) or identified within the signal peptide of MdoD (below). The site of cleavage by signal peptidase is indicated by a white vertical dashed line. **b**, Cryo-EM volume of the *E. coli* TatBC-MdoD complex viewed from the cytoplasm. SP, signal peptide (contour level 0.17). **c**, Model for the MdoD signal peptide shown in the cryo-EM volume (transparent grey surface). **d-i**, Various views of details of the TatBC-MdoD complex with proteins shown as cartoon representations and lipid/detergent densities in white. **d**, One TatBC-signal peptide unit viewed from the membrane plane. **e**, Views from a position in the membrane plane (left) and from within the interior of the complex facing towards the cytoplasm (right). **f**, View from the cytoplasm. **g**, Structural alignment of the apo– and signal peptide-bound –TatBC complexes (arrow shows movements on signal peptide binding). The selected portion of the complex is viewed from the membrane plane (left) or the cytoplasm (right). **h**, TatC Leu-99 is close to lipid/detergent densities within the TatC lipid cavity. **i**, Residues forming the signal peptide-TatB APH interface. **j**,**k,** TatC L99S is required for Tat activity. *E. coli* strains expressing the chromosomal copy of *tatC* with a C-terminal TwinStrep affinity tag and the indicated single amino acid substitutions were analysed. **j**, Tat function was assessed through the ability to grow in the presence of SDS. Strains expressing TatC E103A^44^ or deleted for all *tat* genes (Δ*tatABCDE*) were used as transport-inactive controls. **k**, The ability of the variant TatBC complexes to bind substrate molecules was assessed by asking whether the Tat substrate protein CueO is pulled from solution by the resin-immobilised TatBC complexes. Proteins were detected by immunoblotting. A TatBC(E103A) variant was used as a binding-negative control^40^.

To obtain structural insight into signal peptide recognition by the TatBC complex we collected cryo-EM data from complexes assembled *in vitro* between TatBC and either substrate proteins or signal peptides (Figs. 3-4 and S7-9,11). We then focused classification on the core TatBC region (Fig. S7a-c). The highest resolution volume (2.4 Å) for this region was obtained for a complex between *E. coli* TatBC and three copies of the *E. coli* Tat substrate MdoD (alternative name OpgD) (Figs. 3b and S7a-c). Densities corresponding to three copies of the substrate signal peptide were well resolved at the cytoplasmic face of the trimeric TatBC complex allowing confident model building (Fig. 3b,c). We were also able to resolve a complex between *N. salsuginis* TatBC and the signal peptide of the *E. coli* Tat substrate CueO. Although the signal peptide density in this map was weak, it was consistent with the higher resolution *E. coli* TatBC-MdoD volume (Figs. S8a-f,11).

**Fig. 4:**
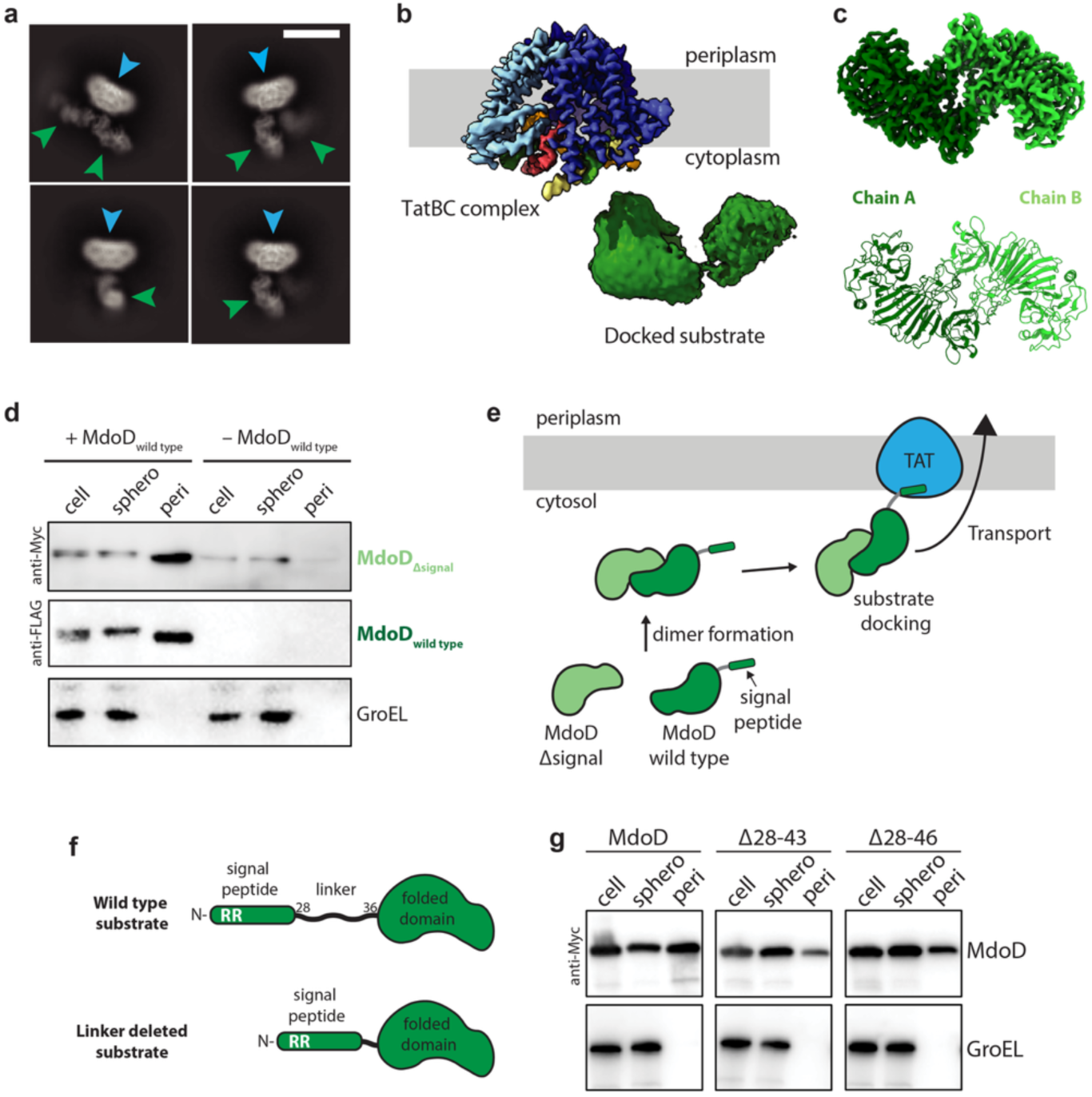
Docking and transport of full-length Tat substrates. **a**, Representative 2D class averages of side views of a TatBC-MdoD complex showing the core TatBC complex (blue arrow) and additional densities corresponding to docked MdoD substrates (green arrow). Scale bar 100 Å. **b**, Cryo-EM volume of the full TatBC-MdoD complex viewed from the membrane plane. Density corresponding to the TatBC complex and to the bound MdoD substrate is shown at a contour level of 1.2 and 0.55, respectively. The membrane, assigned from the position of the detergent micelle, is depicted by a grey box. **c**, Cryo-EM volume at a resolution of 2.1 Å (top) and model (bottom) of the *E. coli* MdoD dimer (contour level 0.24). **d**, A MdoD variant lacking the Tat signal peptide can be exported when co-expressed with signal peptide-bearing MdoD. Transport was assessed in an *E. coli* strain producing a MdoD variant lacking the Tat signal peptide (MdoD_Δsignal_) either with or without co-production of full length MdoD (MdoD_wild type_). The MdoD proteins were epitope tagged with either a myc tag (MdoD_Δsignal_) or FLAG tag (MdoD_wild type_). Whole cells (cell), spheroplast (sphero), and periplasmic (peri) fractions were subjected to immunoblotting with antibodies against the MdoD epitope tags or the cytoplasmic marker protein GroEL. **e**, Schematic showing the process of MdoD_Δsignal_ transport based on the results of 4d. A MdoD dimer formed in the cytoplasm between MdoD_Δsignal_ and MdoD_wild type_ can engage with the Tat export machinery *via* the Tat signal peptide on MdoD_wild type_. **f**, A schematic comparing wild type MdoD (top) with a linker-deleted MdoD variant (bottom). The N-terminal Tat signal peptide of MdoD is connected to the folded domain by a linker region (residues 28-36). **g**, Comparison of the transport competence of wild type MdoD and variants with deletions of amino acids 28 to 43 (Δ28-43) or 28 to 46 (Δ28-46). Whole cells (cell), spheroplast (sphero) and periplasmic (peri) fractions were subjected to immunoblotting to identify MdoD or the cytoplasmic marker protein GroEL.

The TatBC–MdoD complex structure shows that the bound signal peptide is predominantly helical, with a long helix (H1, residues 2-20) encompassing the n– and h-regions and a shorter helix (H2, residues 22-26) corresponding to the c-region (Fig. 3c). When viewed from the cytoplasm, H1 runs from the periphery of the complex towards the centre (Fig. 3b,d-f), and the n-region interacts with the cytoplasmic ends of TMHs 1, 2, and 3 from the same TatC protomer (Figs. 3e, S5b). The APHs from two neighbouring TatB protomers also interact extensively with the signal peptide (Figs. 3e-f, S5b). The h-region threads through the TatB cage, passing over (as viewed from the cytoplasm) one of the interacting TatB APHs. The turn into the c-region then packs under the N-termini of the two signal peptide-interacting copies of TatB in the centre of the complex (Fig. 3e). From there, the c-region helix (H2) exits the central cavity towards the cytoplasm, while maintaining interactions with the APH of the other TatB molecule (Fig. 3d-f). The hydrophobic h-region makes extensive interactions with lipids packed on both sides of this part of the signal peptide (Fig. 3e-i). We observe two structural changes in the TatBC complex upon signal peptide binding. First, a subtle closing of the ends of TatC TMH2 and TMH3 and their connecting loop around the signal peptide n-region (Fig. 3g) driven by bonding interactions with the signal peptide consensus motif (below). Second, the TatB APH becomes more ordered, suggesting a stabilization of its structure in response to signal engagement. (Fig. S8g).

Side chain density for key residues within the signal peptide motif (S-R-R-x-F-L-K) permitted confident modelling (Fig. 3c), allowing us to determine the role of these conserved residues in substrate recognition. The first position in the motif is normally occupied by a residue with high helix-capping propensity(*16*) and our structure confirms that this residue (MdoD D2) caps helix H1 (Fig. 3e,f). The side chain of the first arginine in the motif (MdoD R3) is hydrogen-bonded via an intermediary water molecule to E15 in TatC TMH1, and stacks under F94 from TatC TMH2 (Fig. 3e). The second arginine in the motif (MdoD R4) forms a salt bridge to E103 in TatC TMH3 and packs against Y100 in the TMH2-3 loop (Fig. 3e,f). The consensus phenylalanine (MdoD F6) packs into a hydrophobic pocket formed by F94 in TatC TMH2, and L9, H12, L13 in TatC TMH1 (Fig. 3e). The residues at the consensus leucine and lysine positions (MdoD I7 and K8) interact primarily with lipid densities, with I7 interacting with the hydrophobic tails and K8 positioned to interact with the less well-ordered head groups (Fig. 3e,f). Notably, R4 is also positioned to interact with these lipid head groups (Fig. 3e). The key TatC residues involved in signal peptide recognition (H12, E15, F94, Y100, E103; Fig. 3e) are highly conserved and have previously been implicated in substate binding by biochemical and genetic studies(*21, 24, 25, 48*).

A further TatC residue that has been linked to signal peptide recognition and function is L99(*21, 25, 49, 50*). We now see that this residue is not involved in direct interactions with the signal peptide but rather forms part of the pocket trapping lipids between the Tat core complex and the signal peptide (Fig. 3e-i) Equivalent lipid densities are also present in the signal peptide-bound *N. salsuginis* TatBC map (Extended Data Fig. 7f). Given the earlier data on the functional importance of L99, these structural observations suggest that lipids bound within the core complex contribute to signal peptide recognition. Support for the idea that L99-lipid interactions are important for signal peptide recognition comes from the observation that a Leu to Ser change at this position to increase side chain polarity but not bulk, abolishes not only Tat function (Fig. 3j) but also substrate binding (Fig. 3k).

## Association of full-length substrate proteins with the TatBC complex

Further analysis of our TatBC-MdoD dataset allowed us to examine how the folded domains of substrates are positioned when their signal peptide is engaged at the Tat export machinery. Re-extraction of the particles into a larger box and new 2D classifications generated 2D class averages with density corresponding to the folded passenger domain of MdoD (Fig. 4a, green arrows) positioned proximal to the TatBC complex (Fig. 4a, blue arrows). The majority of 2D classes only showed evidence of one or two relatively well-ordered MdoD passenger domains docked at each TatBC complex (Fig. 4a), even though we had observed full occupancy of the three signal peptide binding sites (Fig. 3b). This implies that the ordering of the folded domain with respect to the TatBC complex is not precise and that the ‘missing’ folded domains reflect high variability in the location of the folded domains rather than their absence.

A cryo-EM map was generated for the complex between TatBC and the most ordered of the bound MdoD proteins (Figs. 4b and S7d). This reveals that the folded domain of the substrate does not form direct, ordered contacts with the membrane portion of the TatBC complex (Fig. 4b). The folded MdoD domain is less well-resolved than the TatBC portion, likely due to mobility of even this most ordered copy of the substrate protein around a flexible tether sequence (below) that links the TatBC-bound signal peptide to the folded domain of the substrate protein (Fig. 4b).

Focused refinement on the substrate density in the TatBC-MdoD complex yielded a 2.1 Å map for MdoD (Figs. 4c and S7d). The map shows that MdoD is a tail-to-tail homodimer, consistent with a recent crystal structure(*51*). Homo-oligomeric Tat substrates, such as MdoD, have a signal peptide on each protomer and thus the potential for each subunit to be independently exported. Given this possibility, it has been a long-standing question whether the Tat system transports homo-oligomeric proteins before or after oligomer formation has taken place(*27*). This question is not resolved by our structural observation that TatBC can bind dimeric MdoD, because our complex was formed by *in vitro* reconstitution rather than through stalling substrate transport *in vivo*. However, we were able to show that MdoD protomers without a signal peptide can be carried to the periplasm by signal peptide-containing protomers (Fig. 4d,e). This demonstrates that MdoD dimer formation must occur in the cytoplasm before Tat transport takes place. It also shows that only one Tat signal peptide on each MdoD dimer is required for transport, a conclusion that is consistent with our structural data showing MdoD positioned so that only one of the signal peptides in the dimer can interact with the TatBC complex (Fig. 4b,e). More generally our MdoD experiments suggest that homo-oligomeric Tat substrates are transported after the constituent subunits have assembled, as is the case for hetero-oligomeric Tat substrates(*2, 52, 53*)

In our TatBC-MdoD complex volumes the first 27 residues of the 32 residue MdoD signal peptide can be modelled before the density weakens (Figs. 3c and S9a). MdoD density is next resolvable at residue 36, though remains poorly ordered and with minimal contacts to the folded part of the molecule until residue 48 (Figs. 4f and S9b). Thus, there is a tether of at least 7, and possibly as many as 19, flexible amino acids linking the bound signal peptide to the folded domain of the substrate (Figs. 4f and S9b). Deletions extending even to the entirety of this flexible tether (residues 28-46) reduce but do not abolish export of the substrate protein (Fig. 4g). This indicates that a stretch of unstructured protein between the signal peptide and folded domain of the substrate protein promotes efficient transport but is not essential for substrate transport to occur.

## TatA introduces asymmetry into the Tat core complex

TatA is structurally similar to TatB and biochemical evidence suggests that it is capable of occupying the TatB site of the Tat core complex when TatB is absent(*35*). It has also recently been proposed that during the Tat translocation cycle TatB is displaced from its binding site by TatA moving from a peripheral site near TatC TMH6(*37, 38*). However, our TatBC complex structures show that the TatB binding site is buried within the core complex and unlikely to be easily exchanged. To clarify how the core complex interacts with TatA we attempted to purify and obtain structural information on TatAC complexes.

An *E. coli* TatAC complex structure could not be reconstructed due to extreme compositional heterogeneity. However, we were able to determine a 3.9 Å resolution reconstruction of the *N. salsuginis* TatAC complex (Figs. 5a and S10,12). In this complex the TatA TMH occupies precisely the same site as the TatB TMH in the TatBC structures (Figs. 5b and S12a-c) and the structure of the TatC components is unaltered within experimental error. The highly conserved polar amino acid at TatA position 8 (E8 in *N. salsuginis* TatA, corresponding to Q8 in *E. coli* TatA) makes equivalent interactions to TatB E8, while the hydrophobic residues in the TatA TMH make analogous packing interactions to TatB with the adjacent TatC molecules (Fig. 5c). Overall, despite sequence differences between the TMHs of TatA and TatB, all TatB TMH interactions with TatC are preserved in the TatAC complex (Fig. 5c). The most striking difference between the TatAC and TatBC structures is the lack of observable density for the TatA APH, suggesting that it is more mobile than the well-ordered APH of TatB (Fig. S12a,b). This interpretation is supported by comparative molecular dynamics simulations of TatAC and TatBC complexes, which show that the TatA APH is highly mobile, when compared with TatB (Fig. S10d-e).

**Fig. 5:**
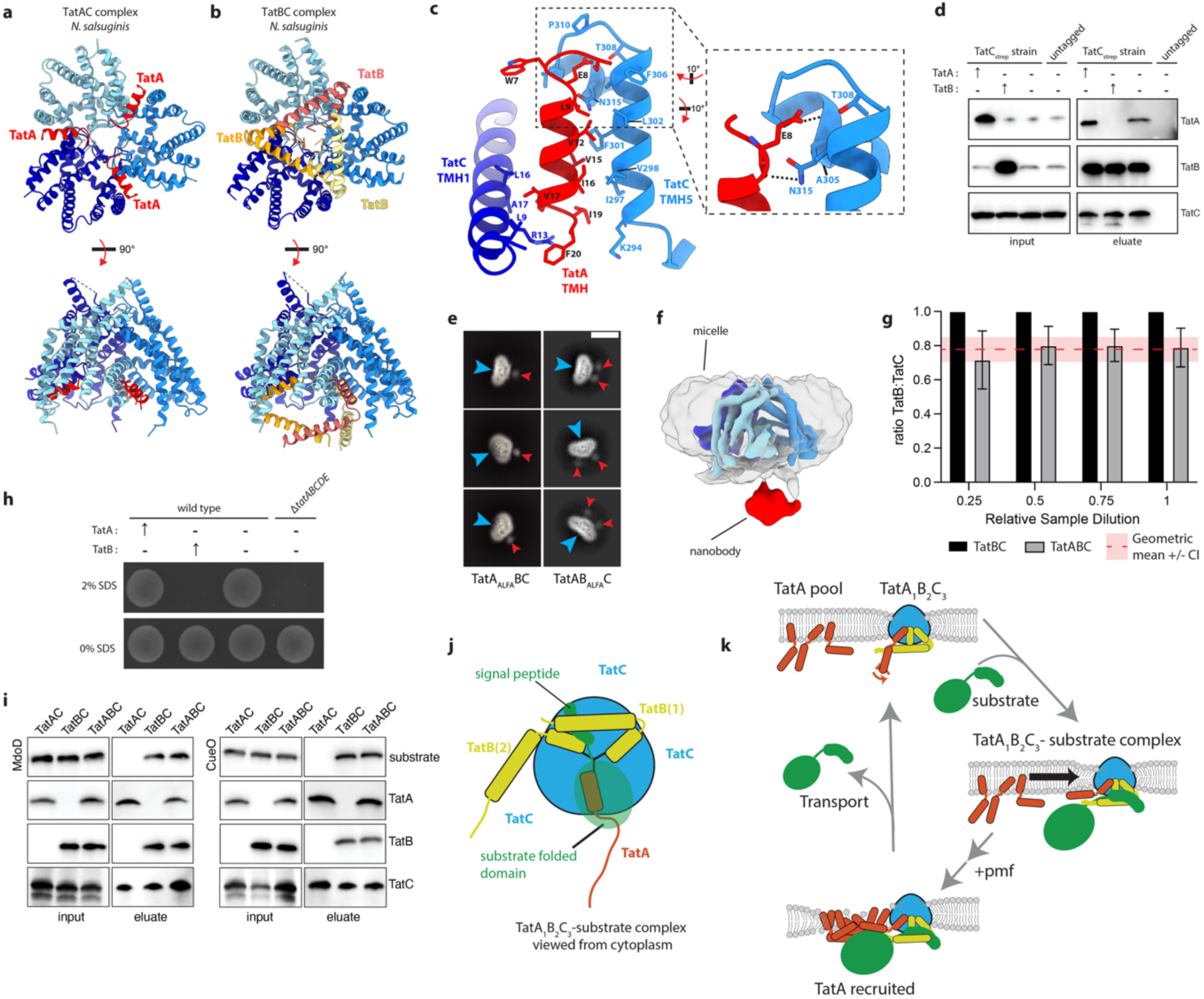
TatA is a constitutive component of the Tat receptor complex. **a**,**b** Models of the *N. salsuginis* (**a**) TatAC and (**b**) TatBC complexes viewed from the cytoplasm (top) and the membrane plane (bottom). TatC subunits are depicted in various shades of blue. **c,** Structure of a *N. salsuginis* TatA-TatC interface showing the interactions between the TatA TMH and TatC-TMH1 and TatC-TMH5. Residues involved in TatA-TatC contacts are labelled. The zoomed in view (inset panel right) highlights the interactions found at the previously proposed ‘polar cluster site’. Hydrogen bonds are depicted by dashed lines. **d**, Demonstration that the native *E. coli* core complex contains TatA and examination of the effects of TatA and TatB overproduction on the loading of TatA and TatB into the complex. Native core complexes isolated through modification of the TatC protein with a Twin-Strep tag and streptactin affinity chromatography are characterized by immunoblotting. The blots characterize the fully native complex (lane 3, both blots) or the complexes present when TatA or TatB are additionally overproduced (↑) from an expression plasmid (lanes 1 and 2, both blots). A wild type (untagged TatC) strain was used as a negative control (lane 4, both blots). **e,** Representative 2D class averages of side views of the *E. coli* TatABC complexes with either TatA (LH panel) or TatB (RH panel)(blue arrows) labelled with a nanobody (red arrows) directed against an inserted ALFA tag in either TatA or TatB respectively. Scale bar 100 Å. **f,** Cryo-EM volume for the nanobody-bound TatABC complex viewed from the membrane plane. Density corresponding to the TatC molecules is coloured in various shades of blue (contour level 0.5), to the surrounding detergent micelle in grey, and to the nanobody in red (both contoured to 0.25). **g**, Ratio of TatB to TatC in TatBC and TatABC samples quantified from Coomassie stained gels at 4 dilutions (total of 7 repeats from three independent samples at each dilution) demonstrates that there is relatively less TatB in the TatABC complex. Geometric mean 0.78, 95% confidence interval 0.71 to 0.85, p=0.003 (ANNOVA, Tukey’s Test). **h**, Effect of TatA and TatB overproduction on Tat transport. TatA or TatB were overproduced where indicated (↑) from a plasmid in either a Tat wild type or Tat null (Δ*tatABCDE*) background and Tat function assessed through the ability to grow in the presence of SDS. **i,** Only core complexes containing TatB are able to bind Tat substrates. *N. salsuginis* TatA_His_C_strep_, TatB_FLAG_C_strep_, or TatA_His_B_FLAG_C_strep_ complexes were immobilised on streptactin affinity resin and incubated with either purified MdoD_Myc_ (left) or CueO_Myc_ (right) Tat substrates. After extensive washing, proteins were eluted and subjected to immunoblotting with antibodies against the protein epitope tags. **j**, Model for substrate binding by the Tat core complex based on the results of this study. The core complex is composed of one copy of TatA, two copies of TatB and three copies of TatC. There is one fully formed substrate binding site (that is one that includes two TatB molecules). The substrate signal peptide binds at this site, positioning the substrate folded domain (shown in transparent rendering) at the opposite side of the complex and adjacent to the TatA site. **k**, Updated model of the Tat translocation mechanism based on the results of this study. A TatA_1_TatB_2_TatC_3_ core complex thins the membrane around the complex while a large pool of TatA molecules is located in the bulk membrane. Substrate docking at the core complex (panel j) triggers pmf-dependent accumulation of additional copies of TatA in the vicinity of the core complex aided by the local membrane thinning, interactions with the substrate folded domain, and changes in the position of the APH of the core complex TatA molecule. Concentrating the TatA molecules results in further local thinning of the membrane bilayer proximal to the substrate folded domain promoting substrate translocation across the membrane bilayer.

We next asked how TatA is incorporated into the full core complex containing all three Tat components. To ensure that we were structurally characterising the physiologically relevant TatABC complex, we initially purified endogenous *E. coli* complexes using an affinity tag on a chromosomally-encoded TatC protein. Whilst this preparation confirmed that native Tat complexes contain TatA in addition to TatB and TatC(*35*) (Fig. 5d) it produced insufficient material for cryo-EM analysis. We, therefore, moved to purifying TatABC complexes from cells expressing the *tatABC* operon from a low copy number plasmid. To assess whether all Tat complexes contain TatA we used a dual-affinity tag strategy with distinct affinity tags on TatA and TatC, purifying sequentially, first by the tag on TatC, followed by the tag on TatA. There was no increase in the TatA-to-TatC ratio from input to eluate after the second purification step targeting TatA (Fig. S12d), indicating that all the complexes isolated via the TatC affinity tag also contain TatA.

Cryo-EM data were collected for TatABC complexes with and without bound substrate. The highest resolution volume obtained was from a TatABC with bound CueO substrate and yielded a 3.3 Å reconstruction (Fig. S13) that was comparable to our TatBC structures. This similarity demonstrates that incorporation of TatA does not induce any fundamental structural rearrangements within the core complex. Notably, no additional density was observed at the periphery of the complex, where TatA has previously been proposed to localize(*37, 38*), suggesting that TatA occupies positions equivalent to TatB. However, the lack of side chain detail on the TMHs did not allow sites occupied by the TatA or TatB TMHs to be distinguished despite extensive classifications and realignments. To visualise TatA in the 2D classes, we inserted an ALFA tag into the cytoplasmic part of TatA to allow TatA protomers within the core complex to be identified through the binding and visualization of an anti-ALFA nanobody (Fig. S14). 2D class averages of the nanobody-bound TatABC complex contained additional density adjacent to the core complex, which we assign to the bound nanobody (Fig. 5e). Weak density corresponding to the ALFA nanobody was also visible in the corresponding 3D reconstructions (Figs. 5f and S14). All particles from which the Tat complex could be reconstructed appeared to have a single bound nanobody (Fig. 5f), indicating that the core complex contains one TatA molecule. By contrast, introduction of an analogous ALFA tag into TatB led to 2D class averages in which two additional densities adjacent to the core complex (Fig. 5e) could be detected suggesting that there are two copies of TatB in this object. From these observations we infer that the core complex has a TatA_1_B_2_C_3_ subunit stoichiometry, and that a TatA molecule replaces one of the three TatB subunits found in the TatBC complex thereby forming an inherently asymmetric object. Consistent with this interpretation we find that the ratio of TatB to TatC is significantly reduced in the TatABC complex relative to the TatBC complex (Fig. 5g). If the three structurally identical TatA/B binding sites in the Tat core complex need to be filled at a fixed stoichiometry of one TatA to two TatB subunits, then the cell would be expected to maintain careful control over the levels of TatA and TatB to ensure appropriate loading of the sites. To test this prediction, we characterized natively expressed core complexes from strains overproducing either TatA or TatB (Fig. 5d). Increasing the cellular levels of TatB markedly reduced the levels of natively expressed TatA that co-purifies with TatC confirming that TatB competes with TatA for assembly into the complex. In contrast, raised levels of TatA did not disrupt TatB incorporation into the complex, likely because TatB has a higher affinity for TatC than TatA. Under native conditions TatB must bind to TatC despite the presence of a large excess of TatA(*7, 54, 55*).

If the stoichiometric inclusion of TatA in the Tat core complex is important for Tat function, then the observed displacement of TatA from the complex by TatB overproduction would be predicted to affect Tat transport. This is indeed the case since we find that overproduction of TatB blocks growth in the presence of SDS which depends on Tat-exported amidases(*56*) (Fig. 5h). By contrast, overproduction of TatA does not impede growth on SDS, consistent with our observation that increasing TatA levels does not affect TatB loading of the core complex. A previous report that overproduction of TatB in an otherwise native Tat background blocks Tat export is also consistent with these observations(*57*). Taken together these data support the hypothesis that incorporation of TatA in the full complex is essential for function and that the functionally relevant Tat core complex is an asymmetric TatA_1_B_2_C_3_ complex.

To further probe the function of TatA relative to TatB in the core complex we compared the ability of purified *N. salsuginis* TatAC, TatBC, and TatABC complexes to form stable complexes with the *E. coli* substrate proteins CueO and MdoD (a well-behaved *E. coli* TatAC complex not being available for this purpose). Only those core complexes containing TatB protomers were able to bind substrate proteins (Fig. 5i). This suggests that the TatA-containing site in the TatABC complex is unlikely to be the primary site of substrate docking. This inference is supported by analysis of the TatB-signal peptide contacts which reveals that only 4 of the 15 interactions are likely able to be recapitulated by the TatA sequence (c.f. all of the TatB-TatC contacts Figs. 5c and S5c) and by MD simulations that show that TatAC complexes coordinate signal peptides less stably that TatBC complexes (Fig. S12f-i). In the TatBC-substrate complexes, substrate binding involves wedging the substrate signal peptide between the APH of TatB and the body of the complex (Fig. 3b,d). However, as discussed above, the APH of TatA is highly mobile in the TatAC complex (Figs. 5a amd S12f-I,14) and may be unable to fulfil an equivalent clamping function. (Fig. S5c) This provides a possible molecular explanation for why only TatB supports tight substrate binding.

## Discussion

Here we provide structural snapshots of the Tat core complex both in the resting state and during the earliest stage of transport where the substrate protein first associates with the core complex. Our structures show that the core complex recognises substrate proteins exclusively through their signal peptides, as previously anticipated(*58*). Binding involves specific protein-protein interactions between residues in the signal peptide n-region consensus motif and residues in TatC. We find that the h-region of the signal peptide is clamped in place by both the TMH and APH of neighbouring copies of TatB, explaining earlier observations that this part of the signal peptide h-region is close to TatB(*26, 59*). Unexpectedly, the signal peptide is also in contact with ordered phospholipids located in the interior of the core complex, at least some of which we show are of functional importance.

The mode of binding of Tat signal peptides to the core complex can be compared and contrasted with the interactions of the structurally similar (though lacking a conserved sequence motif) signal peptides used to target substrates to the Sec translocon. Whilst Tat signal peptides bind to the cytoplasmic face of the core complex (at least in the initial binding state characterized here), Sec signal peptides adopt a membrane-spanning orientation within the Sec apparatus which they are partially exposed to the external membrane bilayer(*60*). However, both types of signal peptide are now seen to have lipid interactions, these being with lipids trapped within the core complex in the case of Tat, but are with the hydrophobic interior of the surrounding bilayer (h-region) and phosopholipid head groups (positively charged n-region) in the case of Sec.

In addition to characterizing the initial core complex-substrate complex, our work uncovers fundamental features of the core complex relevant to later stages in the Tat transport cycle.

First, the structure of the core complex strongly disfavours earlier mechanistic models in which substrate transport is envisaged to occur through the interior of the core complex(*3, 4, 61–65*). Although our structures show that the core complex has a substantial internal cavity, protein transfer through the centre of the complex is highly unlikely for three reasons. First, because this cavity is lined with hydrophobic residues and filled with lipids rather than forming an aqueous environment that folded water-soluble proteins could pass through. Second, because the TatC subunits are tightly packed together at the periplasmic face of the complex and unlikely to open to allow substrate egress (and in the *M. xanthus* complex would additionally be blocked by the highly structured periplasmic domain, Fig. 1c), an inference that is supported by the stability of the complex in MD simulations (Fig. 2e). Third, because the cavity within the complex is not large enough to accommodate the folded domain of many Tat substrates (Fig. S15). Thus, our structural data suggest that substrate transport must occur on the periphery of the core complex(*22, 35*).

Second, the transmembrane regions of the Tat core complex are highly tilted leading to extreme thinning of the surrounding membrane bilayer. This bilayer thinning likely aids the recruitment of TatA to the core complex during the later stages of the translocation cycle. Recruitment of TatA in the thinned membrane around the complex will be favoured in order to relieve the hydrophobic mismatch between the short TatA TMH and the width of the bulk membrane(*31*). This inference is consistent with substrate transport occurring on the periphery of the core complex (above) and supports mechanistic models that incorporate membrane thinning as critical for transport(*31, 43, 66–69*). Emerging evidence indicates that membrane thinning is a common feature of protein translocation and membrane protein insertion systems(*70*).

Third, we find that the physiological core complex contains an unequal number of TatA and TatB subunits each occupying an equivalent site between the TatC subunits, with our data indicating that the stoichiometry is likely TatA_1_TatB_2_TatC_3_. The result is that the physiological core complex is inherently asymmetric. Since our biochemical studies indicate only TatB and not TatA is able to support substrate binding to the core complex, and our substrate-bound structures show that two copies of TatB are needed to create one signal peptide-binding site, we infer that there is only a single physiologically relevant signal peptide-binding site in a TatA_1_B_2_C_3_ complex (there is only one TatC subunit for which signal peptide binding is assisted by two copies of TatB) (Fig. 5i). This TatC subunit is positioned across the complex from the TatA molecule (Fig. 5i). Consequently, the orientation of the bound signal peptide across the centre of the core complex that we observe in this work will direct the attached substrate folded domain towards the TatA site (Fig. 5i). Thus, in this model the asymmetry of the core complex establishes a mode of substrate binding in which the folded domain is positioned for transport at the TatA side of the core complex (Fig. 5i). We see no substantive structural change in the core complex upon substrate binding that could act as a trigger for TatA recruitment. Consequently, we infer that the bound substrate molecule itself clusters TatA molecules for transport though semi-specific interactions with the folded passenger domain.

## Methods

### Bacterial strains and plasmids

Bacterial strains and plasmids used in this study are listed in Table S3. The genes encoding *Myxococcus xanthus* TatB and TatC, and *Nitratifractor salsuginis* TatA, TatB and TatC were codon-optimized for expression in *Escherichia coli* and synthesized as DNA fragments containing homologous recombination sites for Gibson assembly. PCR fragments for *E. coli tatA*, *tatB*, *tatC*, *cueO* and *mdoD* were generated by PCR using Q5 polymerase (NEB) and *E. coli* genomic DNA (DSM 1116). Gibson assembly and PCR reactions were carried out according to the manufacturer’s recommendations.

### Chromosomal modifications

Gene deletions, point mutations and the insertion of sequence encoding strep tags in the bacterial chromosome were performed using the pWRG730 and pWRG717 constructs, using a modified protocol based on Hoffmann *et al*., 2017(*71*). Briefly, an aphI-I-SceI kanamycin resistance cassette was amplified from pWRG717 with primers that incorporate homology regions to the target gene. *E. coli* strains carrying the temperature sensitive construct pWRG730 were grown at 30°C to A_600nm_ = 0.4-0.6. λ red recombinase expression was induced by incubating cultures in a 42°C water bath for 13 minutes, cultures were subsequently incubated on ice for 20 minutes. Cells were harvested by centrifugation and washed three times in ice-cold water. Cells were resuspended in 100 μl of ice-cold water and 100 ng of PCR product was added. Cells were electroporated and recovered in 0.5 ml of SOC media for 1 hour at 30°C. Colonies were selected on LB-agar containing kanamycin (50 μg/ml) and chloramphenicol (10 μg/ml). To introduce point mutations or tags onto the chromosome, a second PCR product was generated that incorporated the point mutation or tag and at least 400 bp flanking the gene of interest. λ red recombinase expression in strains carrying the aphI-I-SceI kanamycin resistance cassette was carried as described above and electroporated with 500 – 1000 ng of the PCR product containing the point mutations or tag. Cells were selected on LB-agar plates containing chloramphenicol (10 μg/ml) and anhydrotetracycline (500 ng/μl). Positive clones were verified by PCR and DNA sequencing.

### Purification of *M. xanthus* and *N. salsuginis* TatBC, and TatAC complexes

*M. xanthus* and *N. salsuginis* TatBC complexes or the *N. salsuginis* TatAC complex were overexpressed in *E. coli* strain L56 carrying both pTatC-GFP-His (pWALDO based) and pTatB-strep (pCDFDuet-1 based) or pTatAstrep (pCDFDuet-1 based). Cells were grown in terrific broth (TB) media containing spectinomycin (100 μg/ml) and kanamycin (50 μg/ml) at 37°C with shaking to A_600nm_ = 4.0 after which cultures were supplemented with 0.1 mM isopropyl ß-D-1-thiogalactopyranoside (IPTG) and incubated for an additional 12 h at 25°C with shaking. Cells were harvested by centrifugation (5,000*g,* 10 min). Purification steps were similar for *M. xanthus* and *N. salsuginus* constructs and carried out at 4°C. Briefly, cell pellets were resuspended in PBS (10 mM Na_2_HPO_4_, 1.8 mM KH_2_PO_4_, 2.7 mM KCl, 137 mM NaCl) supplemented with DNase I (30 μg/ml), lysozyme (500 μg/ml) and a cOmplete EDTA-free Protease Inhibitor Cocktail tablet (Sigma) for 30 minutes before passing through an EmulsiFlex C5 homogenizer (Avestin) at 15,000 psi. Lysates were clarified by centrifugation (30,000*g*, 30 min) and membranes were collected by centrifugation (200,000*g* for 1.5 h). Membranes were resuspended in buffer A (50 mM sodium phosphate pH 8.0, 300 mM NaCl) containing 20 mM imidazole and solubilised by incubation with 1% (w/v) glyco-diosgenin (GDN; Anatrace) for 2 h. Insoluble material was removed by centrifugation (100,000*g*, 30 min) and solubilised membranes applied to three 5 ml Ni-NTA superflow cartridges (QIAGEN). The resin was washed with 10 column volumes (10 CV) of buffer A containing 0.02% (w/v) GDN and 20 mM imidazole, followed by washing with 10 CV of buffer A containing 0.02% (w/v) GDN and 40 mM imidazole. Proteins were eluted in 4 CV of buffer A containing 0.02% (w/v) GDN and 300 mM imidazole. TEV protease was added to the eluates at a ratio of TEV to TatBC or TatAC of 1:100 and the sample dialysed overnight using 10,000 molecular weight cutoff (MWCO) SnakeSkin dialysis tubing (ThermoScientific) into buffer B (50 mM sodium phosphate 8.0, 150 mM NaCl, 0.02% GDN). The dialysed sample was adjusted to contain 20 mM imidazole and passed through a 5 ml Ni-NTA superflow cartridge (QIAGEN) to remove GFP-His. The sample was then applied to a 5 ml Streptactin XT superflow column (IBA Lifesciences). The resin was washed with 20 CV of buffer B (50 mM sodium phosphate pH 8.0, 150 mM NaCl, 0.5 mM EDTA) containing 0.02% GDN and proteins eluted in 5 CV of buffer B supplemented with 50 mM D-biotin (IBA Lifesciences) and 0.02% GDN. Eluates were concentrated using a 100 kDa MWCO Vivaspin 20 (cytiva) centrifugal filter unit and injected onto a Superose 6 Increase 10/300 GL size exclusion column (cytiva) pre-equilibrated in buffer C (20 mM HEPES pH 7.4, 150 mM NaCl, 0.02% GDN). Peak fractions were collected and concentrated using a 100 kDa MWCO Vivaspin (GE healthcare) centrifugal filter unit.

### Purification of *E. coli* TatBC and TatABC complexes

*E. coli* TatBC and TatABC complexes were overexpressed in an *E. coli* MC4100 Δ*tatABCD*, Δ*tatE* carrying pQE60 based vectors and pREP4 (listed in Table S3). Cultures were grown in LB media supplemented with 4% glycerol, kanamycin (50 μg/ml) plus carbenicillin (100 μg/ml) or chloramphenicol (25 ug/ml) at 37°C with shaking to A_600nm_ = 0.7. Cells were then supplemented with 1 mM IPTG and incubated for an additional 4 h at 37°C with shaking. Cell pellets were resuspended in lysis buffer (50 mM Tris pH 8.0, 200 mM NaCl, 1 mM EDTA) supplemented with DNase I (30 μg/ml), lysozyme (500 μg/ml) and a cOmplete EDTA-free Protease Inhibitor Cocktail tablet (Sigma) for 30 minutes before passing through an EmulsiFlex C5 homogenizer (Avestin) at 15,000 psi. Lysates were clarified by centrifugation (12,000*g*, 10 min) and membranes were collected by centrifugation (150,000*g*, 1.5 h). Membranes were resuspended in lysis buffer and solubilised by incubation with 1% (w/v) glyco-diosgenin (GDN; Anatrace) for 16 h at 4°C. Insoluble material was removed by centrifugation (150,000*g*, 40 min) and solubilised membranes were supplemented with 12 mM imidazole then mixed with pre-equilibrated cOmplete His-tag purification resin (Roche) for 3 h at 4°C. The resin was centrifuged at 500*g* for 5 min and then transferred to a gravity column. The resin was washed with 15 CV of lysis buffer supplemented with 12 mM imidazole and 0.02 % GDN, followed by elution in lysis buffer supplemented with 100 mM imidazole and 0.02 % GDN. Eluates were concentrated using a 100 kDa MWCO Vivaspin 20 (cytiva) centrifugal filter unit and injected onto a Superose 6 Increase 10/300 GL size exclusion column (cytiva) pre-equilibrated in lysis buffer plus 0.02 % GDN. Peak fractions were collected and concentrated using a 100 kDa MWCO Vivaspin (GE healthcare) centrifugal filter unit.

Membranes from cells overexpressing *E. coli* TatABC complexes containing an internal His tag or ALFA-His tag in TatA or TatB and a C-terminal strep tag in TatC were prepared and solubilised with GDN as described above for the *E. coli* TatBC complex. Insoluble material was removed by centrifugation (100,000*g*, 30 min) and solubilised membranes applied to a 5 ml Streptactin XT superflow column (IBA Lifesciences). The resin was washed with 20 CV of buffer B (50 mM sodium phosphate pH 8.0, 150 mM NaCl, 0.5 mM EDTA) containing 0.02% GDN and proteins eluted in 5 CV of buffer B supplemented with 50 mM D-biotin (IBA Lifesciences) and 0.02% GDN. The sample was adjusted to containing 3 mM imidazole and passed through a cOmplete His-tag purification column (Roche). The column was washed with 20 column volumes (20 CV) of buffer A containing 0.02% (w/v) GDN and 2 mM imidazole. Proteins were eluted in 6 CV of buffer A containing 0.02% (w/v) GDN and 200 mM imidazole. Eluates were concentrated using a 100 kDa MWCO Vivaspin 20 (cytiva) centrifugal filter unit and injected onto a Superose 6 Increase 10/300 GL size exclusion column (cytiva) pre-equilibrated in buffer C (20 mM HEPES pH 7.4, 150 mM NaCl, 0.02% GDN). Peak fractions were collected and concentrated using a 100 kDa MWCO Vivaspin (GE healthcare) centrifugal filter unit.

### Purification of selenomethionine-labelled *M. xanthus* TatC periplasmic domain

The periplasmic domain (incorporating residues 127-303) of *M. xanthus tatC* was subcloned into a pET15b vector by PCR amplification of codon-optimized *M. xanthus tatC*, followed by restriction digestion and ligation according to manufacturer’s recommendations. Protein was overexpressed in the auxotrophic *E. coli* strain B834 (DE3) grown in SelenoMethionine Medium (Molecular Dimensions) plus 100 μg/ml carbenicillin after induction of A_600nm_ = 0.6 cultures with 1 mM IPTG for 18 h at 21°C with shaking. Cells were harvested by centrifugation (5,000*g,* 10 min). Cell pellets were resuspended in lysis buffer containing 20 mM HEPES pH 7.5, 300 mM NaCl, 10 mM imidazole, 2 mM TCEP, and supplemented with DNase I (30 μg/ml), lysozyme (500 μg/ml) and a cOmplete EDTA-free Protease Inhibitor Cocktail tablet (Sigma) for 30 minutes before passing through an EmulsiFlex C5 homogenizer (Avestin) at 15,000 psi. Lysates were clarified by centrifugation (30,000*g*, 30 min) and supernatant was applied to a 5 ml Ni-NTA superflow cartridge (Qiagen) pre-equilibrated with lysis buffer. The resin was washed with 10 CV of lysis buffer followed by 20 CV of lysis buffer supplemented with 20 mM imidazole. Protein was eluted in lysis buffer supplemented with 250 mM imidazole followed by addition of TEV protease (1:100 ratio of TEV to TatC periplasmic domain) and the sample was then dialysed overnight using 3,000 molecular weight cutoff (MWCO) SnakeSkin dialysis tubing (ThermoScientific) into 20 mM HEPES pH 7.5, 150 mM NaCl, 2 mM TCEP. The dialysed sample was adjusted to contain 20 mM imidazole and passed through a 5 ml Ni-NTA superflow cartridge (QIAGEN) to separate the proteolyzed His_6_-tag and His-tagged TEV protease from the flow-through. The flow-through was subsequently concentrated using a 10 kDa MWCO Vivaspin 20 (cytiva) centrifugal filter and injected onto a Superdex 75 26/600 size exclusion column (cytiva) pre-equilibrated in buffer containing 20 mM HEPES pH 7.5, 150 mM NaCl, 2 mM TCEP. Peak fractions were collected and concentrated to an A_280nm_ of 8.9 using a 10 kDa MWCO Vivaspin (cytiva) centrifugal filter unit for crystallization.

### Crystallization and data collection and processing

Crystallization was carried out using the sitting-drop vapour diffusion method at 21 °C. Crystals were obtained in 0.2 M lithium sulphate, 0.1 M Tris pH 8.5, 10% PEG 8000, 10% PEG 1000 at a 1:1 ratio of protein:mother liquor and a drop volume of 400 nl. These were flash frozen in liquid nitrogen in the mother liquor supplemented with 25% ethylene glycol. Diffraction data were collected to 2.0 Å at the Diamond light source (Beamline I02), at 120K from 1 SeMet-labelled crystal (λ = 0.97926). Data were indexed, integrated and scaled using iMosflm(*72*) and point and scale within CCP4. Eighteen SeMet sites were detected using Phenix Autosol(*73*) with a FOM of 0.327. The data were phased, density modified and initial model built using Phenix Resolve(*74*) (map skew 0.3, correlation of local RMS density 0.87). The model was iteratively manually adjusted in Coot(*75*) and refined in Phenix(*76*) to produce the model described in Table S2.

### Purification of TatABC– and TatBC-substrate complexes

Full length MdoD or CueO was expressed in *E. coli* strain BL21 (DE3) Δ*tatABCD*, Δ*tatE* from a pET28a vector modified to encode an N-terminal His_6_-SUMO fusion. Cells were grown in LB media containing kanamycin (50 μg/ml) at 37°C with shaking to A_600nm_ = 0.8 after which cultures were supplemented with 1 mM IPTG and incubated for an additional 12 h at 18°C with shaking. Cells were harvested by centrifugation (5,000*g,* 10 min). Cell pellets were resuspended in Buffer D (50 mM Tris.HCl pH 8.0, 500 mM NaCl, 1 mM EDTA and 2 mM imidazole) supplemented with DNase I (30 μg/ml), lysozyme (500 μg/ml) and a cOmplete EDTA-free Protease Inhibitor Cocktail tablet (Sigma) for 30 minutes before passing through an EmulsiFlex C5 homogenizer (Avestin) at 15,000 psi. Lysates were clarified by centrifugation (30,000*g*, 40 min) and applied to a 1 ml cOmplete His-tag purification column (Roche) and the resin washed with 40 CV of buffer D. Proteins were eluted in 6 CV of buffer E (50 mM Tris.HCl pH 8.0, 150 mM NaCl, 1 mM EDTA and 200 mM imidazole) and the eluate dialysed into 20 mM HEPES pH 7.4, 150 mM NaCl, 1 mM EDTA using 10,000 molecular weight cutoff (MWCO) SnakeSkin dialysis tubing (ThermoScientific). To assemble the TatBC-MdoD complexe and TatABC-CueO, His_6_-SUMO-MdoD or His_6_-SUMO-CueO was incubated at 4°C for 1 h with *Saccharomyces cerevisiae* Ubiquitin-like-specific protease 1 (Ulp1), the buffer adjusted to contain 0.02% GDN and the samples mixed with purified *E. coli* TatBC or TatABC complex. The samples were incubated overnight at 4°C and injected onto a Superose 6 Increase 10/300 GL size exclusion column (cytiva) pre-equilibrated in 50 mM Tris pH 8.0, 200 mM NaCl, 1 mM EDTA, 0.02% GDN. Peak fractions were collected and concentrated using a 100 kDa MWCO Vivaspin (GE healthcare) centrifugal filter unit. To assemble the *N. salsuginis* TatBC–CueO signal peptide complex, 50 μl TatBC (A_280nm_ = 8.0) was mixed with 50 μl CueO peptide (1 mg/ml, Peptide Protein Research Ltd., residues 1-28) and incubated for 12 h at 4°C before grid preparation.

### Reconstitution of *N. salsuginis* TatBC into MSPE3D1 nanodiscs

Chloroform was removed from *E. coli* polar lipids (Avanti) using a rotary evaporator. The lipids were washed twice with pentane and then resuspended at 20 mg/ml in MSP buffer (20 mM Tris-HCl pH 7.4, 100 mM NaCl, 1 mM EDTA) containing 60 mM cholate and stored at –80°C until required. *N. salsuginis* TatBC was purified as described above except membranes were solubilised in 1% digitonin instead of GDN and purification buffers contained 0.1% digitonin rather than 0.02% GDN. For nanodisc reconstitution digitonin purified TatBC was mixed with membrane scaffold protein (MSP1E3D1, purified as previously described(*77*)), and *E. coli* polar lipids using a 1:2:122 (TatBC:MSP1E3D1:lipid mixture) molar ratio and diluted with MSP buffer to a final cholate concentration of 25 mM. The reaction mixture was incubated on ice for 1 h, followed by detergent removal using 1 g of pre-washed Biobeads SM-2 (BioRad) per ml of reaction mixture by incubation overnight at 4°C. To enrich for nanodiscs containing TatBC, the nanodisc reaction mixture was applied to a 1 ml Streptactin XT superflow column (IBA Lifesciences), the resin washed with 20 CV of MSP buffer and proteins eluted with MSP buffer supplemented with 50 mM D-biotin (IBA Lifesciences). The eluate was concentrated using a 100 kDa MWCO Vivaspin 20 (cytvia) centrifugal filter unit and injected onto a Superose 6 Increase 10/300 GL size exclusion column (cytvia) pre-equilibrated in MSP buffer. Peak fractions were collected and concentrated using a 100 kDa MWCO Vivaspin (GE healthcare) centrifugal filter unit.

### Cryo-EM sample preparation and imaging

Purified complexes (4 μl each) of *E. coli* TatBC (A_280nm_ = 4.2), *M. xanthus* TatBC (A_280nm_ = 4.4), *N. salsuginis* TatBC (A_280nm_ = 4.0), *N. salsuginis* TatBC in nanodisc (A_280nm_ = 2.5), *N. salsuginis* TatBC-CueO peptide (A_280nm_ = 4.0), *E. coli* TatABC-CueO (A_280nm_ = 4.3), *E. coli* TatBC-MdoD (A_280nm_ = 5.0), *E. coli* TatA_ALFA_BC-ALFA nanobody complex or *N. salsuginis* TatAC (A_280nm_ = 4.0), *E. coli* TatAB_ALFA_C-ALFA nanobody complex (A_280nm_ = 5.5), were adsorbed to glow discharged holey carbon-coated grids (Quantifoil 300 mesh, Au R1.2/1.3 for 15 s at 15 mA). Grids were then blotted for 2 s at 100% humidity at 4°C and frozen in liquid ethane using a Vitrobot Mark IV (FEI). Data were collected in counted mode on a Titan Krios G3 (FEI) operating at 300 kV with a GIF energy filter (Gatan) with slit width of 20 eV at 105,000x magnification and K3 Summit detector (Gatan) or on a CFEG-equipped Titan Krios G4 (Thermo Scientific) operating at 300 kV with a Selectris X imaging filter (Thermo Fisher Scientific) with slit width of 10 eV at 165,000x magnification on a Falcon 4 direct detection camera (Thermo Fisher Scientific). Pixel sizes ranged from 0.693 Å – 0.832 Å and movies were collected at a total dose ranging from 51.1 e^-^/A^2^ – 62.9 e^-^/A^2^ fractionated to 0.9 – 1.5 e^-^ / A^2^ / fraction for motion correction across the various datasets (Table S1).

### Cryo-EM data processing

Patched motion correction, CTF parameter estimation, particle picking, extraction, and initial 2D classification were performed in SIMPLE 3.0(*78*). All downstream processing was carried out in cryoSPARC(*79*) or RELION(*80*),using the csparc2star.py script within UCSF pyem(*81*) to convert between formats. Global resolution was estimated from gold-standard Fourier shell correlations (FSCs) using the 0.143 criterion and local resolution estimation was calculated within cryoSPARC using an FSC threshold of 0.5 or within RELION(*80*).

The cryo-EM processing workflow for *E. coli* TatBC in GDN is outlined in Fig. S2a-c. Briefly, particles were subjected to two rounds of reference-free 2D classification (k=300) using a 160 Å soft circular mask within cryoSPARC. Four volumes were then generated from a 393,435 particle subset of the 2D-cleaned particles after multi-class *ab initio* reconstruction (C3 symmetry) using a maximum resolution cutoff of 6 Å. Output volumes were lowpass-filtered to 8 Å and used as references for a 4-class heterogeneous refinement (C3 symmetry) against the full 2D-cleaned particle set (1,218,611 particles). Particles (468,840) from the most populated and structured class were selected and non-uniform refined, applying C3 symmetry, against their corresponding volume lowpass-filtered to 15 Å, generating a 3.2 Å map. Bayesian polishing was performed in RELION followed by local and global CTF-refinement (fitting beamtilt and trefoil) and non-uniform refinement in cryoSPARC to yield a 3.1 Å map that was used for model building.

The cryo-EM processing workflow for *N. salsuginis* TatBC in GDN is outlined in Fig. S2d-f. Briefly, particles were subjected to two rounds of reference-free 2D classification (k=200) using a 150 Å soft circular mask within cryoSPARC. Selected particles (1,169,012) were then input into multi-class *ab initio* reconstruction (C1 symmetry) using a maximum resolution cutoff of 5 Å, generating four volumes. Particles (471,101) belonging to the most populated and structured class were selected and non-uniform refined, applying C3 symmetry, against their corresponding volume lowpass-filtered to 8 Å, generating a 3.0 Å map. Bayesian polishing was performed in RELION followed an additional round of 2D classification in cryoSPARC (150 Å circular mask, k =200) whereby 371,162 particles were kept for further processing. These particles were non-uniform refined (C3 symmetry) against an 8 Å lowpass-filtered reference, generating a 2.6 Å volume that was further improved to 2.5 Å after local and global CTF-refinement (fitting beamtilt and trefoil).

The cryo-EM processing workflow for *M. xanthus* TatBC in GDN is outlined in Fig. S3. Briefly, particles were subjected to one round of reference-free 2D classification (k=300) using a 150 Å soft circular mask within cryoSPARC. Three volumes were then generated from a 33,176 particle subset of the 2D-cleaned particles after multi-class ab initio reconstruction (C1 symmetry) using a maximum resolution cutoff of 5 Å. Output volumes were lowpass-filtered to 8 Å and used as references for a 3-class heterogeneous refinement (C1 symmetry) against the full 2D-cleaned particle set (1,271,712 particles). Particles (636,367) from the most populated and structured class were selected and non-uniform refined, applying C3 symmetry, against their corresponding volume lowpass-filtered to 8 Å, generating a 3.3 Å map. Bayesian polishing was performed in RELION followed an additional round of 2D classification in cryoSPARC (150 Å circular mask, k =200). 2D-selected particles (530,874) were subjected to non-uniform refinement with C3 symmetry against the pre-polished volume lowpass-filtered to 25 Å, yielding a 3.0 Å reconstruction. Focused alignment-free 3D classification (k = 8, T = 4, 25 iterations) using a soft mask covering TatBC transmembrane helices (TMHs) was then performed, yielding two classes with improved density for the TMH of TatB. Particles belonging to both classes were combined and non-uniform refined (C3 symmetry) against a 15 Å lowpass-filtered reference, followed by local and global CTF-refinement (fitting beamtilt and trefoil), to yield a 3.2 Å map that was used for model building.

The cryo-EM processing workflow for *N. salsuginis* TatBC nanodisc complex is outlined in Fig. S6. 10,945,816 particles were subjected to reference-free 2D classification in cryoSPARC (300 classes). Selected particles (1,114,288) were used to generate an *ab initio* model. This model was low pass filtered to 35 Å and used as a reference for a non-uniform refinement using an initial low pass filter of 8 Å, to yield a 3.7 Å map. Non-uniform refinement in cryoSPARC after Bayesian polishing in RELION generated a final map with a global resolution of 3.1 Å.

The cryo-EM processing workflow for *E. coli* TatBC-MdoD in GDN is outlined in Fig. S7a-c. Two datasets were collected for this sample. For dataset 1, particles were subjected to three rounds of reference-free 2D classification (k=300) using a 180 Å soft circular mask for the first two rounds. Five volumes were then generated from a 138,750 particle subset of the 2D-cleaned particles after multi-class *ab initio* reconstruction (C1 symmetry) and using a maximum resolution cutoff of 7 Å. Output volumes were lowpass-filtered to 8 Å and used as references for a 5-class heterogeneous refinement (C1 symmetry) against the full 2D-cleaned particle set (1,796,522 particles). Particles (350,493) belonging to the class that demonstrated structured TMHs were selected and non-uniform refined, applying C1 symmetry, against their corresponding volume lowpass-filtered to 15 Å, generating a 3.4 Å map. Non-uniform refinement using C3 symmetry further improved map resolution to 3.0 Å. For dataset 2, particles were subjected to two rounds of reference-free 2D classification (k=300) using a 180 Å soft circular mask for the first classification job. 2D-cleaned particles (2,834,346) were then subjected to heterogeneous refinement (C1 symmetry) against the same 8 Å lowpass-filtered references generated from *ab initio* reconstructions described above for dataset 1. Particles (850,952) belonging to the class that demonstrated structured TMHs were selected and non-uniform refined, applying C3 symmetry, against their corresponding volume lowpass-filtered to 15 Å, generating a 2.8 Å map. These particles, along with the 350,493 particles that were curated from dataset 1, were polished in RELION, combined, and non-uniform refined (C3 symmetry) against a 15Å lowpass-filtered volume to generate the 2.4 Å volume used in model building. Local and global CTF-refinement was performed but did not appreciably improve map quality. While the input material for this dataset contained purified TatABC with bound MdoD, we were unable to identify density corresponding to TatA in any of our reconstructions, even after extensive classification schemes targeting the signal peptide, substrate, and subunits of the Tat complex. We therefore refer to this complex as TatBC-MdoD throughout.

The cryo-EM processing workflow for the *E. coli* MdoD substrate is outlined in Fig. 7d-f. The *E. coli* MdoD volume was generated by selecting 2D class averages that corresponded to the *E. coli* mdoD substrate in 2D classifications from datasets 1 and 2 of the *E. coli* TatBC-MdoD in GDN sample. Selected 2D class averages from dataset 1 and dataset 2 were used as input for two separate multi-class *ab initio* reconstructions (C1 symmetry) and using a maximum resolution cutoff of 7 Å. Particles from selected classes from dataset 1 were non-uniform refined, applying C2 symmetry, using a volume lowpass-filtered to 15 Å, generating a 2.8 Å volume. The same procedure was particles from dataset 2 which also generated a 2.8 Å volume. Volumes generated from non-uniform refinements applying C1 symmetry were identical to the C2 volumes. Particles were polished in RELION, combined, and non-uniform refined (C2 symmetry) against a 15Å lowpass-filtered volume to generate the 2.3 Å volume. Local and global CTF-refinement was performed, followed by a non-uniform refinement which generated a 2.1 Å volume used in model building.

The cryo-EM processing workflow for the *N. salsuginis* TatBC-CueO peptide complex is outlined in Fig. S11. 8,504,721 particles were subjected to reference-free 2D classification in cryoSPARC (300 classes). Selected particles (1,426,112) were subjected to a multi-class heterorefinement in cryoSPARC using four 8 Å low-pass filtered *ab initio* models generated from a previous data collection of a TatBC-CueO peptide sample as input, which generated a map with a resolution of 4.2 Å. A 2D classification and a non-uniform refinement in cryoSPARC after Bayesian polishing in RELION improved map quality and generated a final map with a global resolution of 3.2 Å. A local CTF refinement followed by a non-uniform refinement generated a map with a resolution of 3.1 Å. Focused 3D classification without alignment was performed in RELION using a soft mask encompassing the peptide binding cavity from which one highly occupied class (84,834 particles) was selected. This particle set was subjected to non-uniform refinement in cryoSPARC to generate a 3.2 Å volume with strong CueO peptide density.

The cryo-EM processing workflow for the *N. salsuginis* TatAC complex is outlined in Fig. S10. 12,432,717 particles were subjected to reference-free 2D classification in cryoSPARC (k=300). Selected particles (591,032) were used to generate multi-class *ab initio* models which were low pass filtered to 35 Å. The particles from the selected model were used as input for a second multi-class *ab initio* reconstruction which yielded a class containing 93,589 particles. This model was low pass filtered to 35 Å and used as a reference for a non-uniform refinement using an initial low pass filter of 8 Å, to yield a 3.8 Å map. A 2D classification and non-uniform refinement in cryoSPARC after Bayesian polishing in RELION generated a final map with a global resolution of 3.4 Å.

The cryo-EM processing workflow for the *E. coli* TatABC-CueO complex is outlined in Fig. S13. Briefly, 5,301,381 particles were subjected to two rounds of reference-free 2D classification in cryoSPARC (k=200). Five volumes were then generated from 855,150 2D-cleaned particles after multi-class *ab initio* reconstruction (C1 symmetry) using a maximum resolution cutoff of 7 Å. Output volumes were lowpass-filtered to 7 Å and used as references for a 5-class heterogeneous refinement (C1 symmetry) against the same 855,150 2D-cleaned particles. Particles (247,806) from the most populated and structured class were selected and non-uniform refined in C1 against their corresponding volume lowpass-filtered to 15 Å, resulting in a 3.3 Å volume. Extensive classifications schemes were performed on the subunits of the Tat complex and/or signal peptide, but we were unable to resolve any inherent asymmetry.

The cryo-EM processing workflow for the *E. coli* TatA_ALFA_BC – ALFA nanobody complex is outlined in Fig. S3. 6,808,766 particles were subjected to reference-free 2D classification in cryoSPARC (k=200). Selected particles (3,647,437) were subjected to a multi-class heterorefinement in cryoSPARC using four 8 Å low-pass filtered *ab initio* models generated from a subset of particles as input. The particles from the selected class were used as input for another round of 2D classification, selected particles were used as input for two additonal multi-class *ab initio* reconstructions which yielded a class containing 113,148 particles. This model was low pass filtered to 35 Å and used as a reference for a non-uniform refinement using an initial low pass filter of 8 Å, to yield a 6.4 Å map.

Selected 2D class averages of the *E. coli* TatAB_ALFA_C – ALFA nanobody complex were obtained following two rounds of 2D classification (k=200) of a 7,448,890 particle set, a multi-class *ab initio* (k=4) reconstruction to further remove junk particles, and a subsequent 2D classification (k=100) on 303,279 selected particles.

### Cryo-EM model building and refinement

Atomic models were built into their respective cryo-EM volumes in Coot(*75*). Models were further refined in real-space using PHENIX(*76*) with rotamer, Ramachandran restraints, and secondary structure restraints (where necessary) against either global B-factor sharpened maps or deepEMhancer maps, yielding the models described in Table S1. All models were validated using MolProbity within PHENIX(*82*). A homology model of *E. coli* MdoD was generated by sequence threading against the *E. coli* MdoG model using Phyre2(*83*). Figs. were prepared using UCSF ChimeraX(*84*) and Adobe Illustrator.

### SDS sensitivity assay

*E. coli* strains were grown aerobically overnight at 37°C in Luria-Bertani (LB) broth. Cultures were diluted in fresh LB broth grown to A_600nm_ = 1.0. For strains carrying pQE-based vectors, media and plates were supplemented with carbenicillin (100 μg/ml) and IPTG (100 μM – 1 mM). Cultures were diluted 1000-fold in PBS and 5 μl spotted onto LB agar supplemented with 2% SDS and incubated overnight at 37°C. The data presented are representative of at least three independent experiments.

### Substrate export assays

Overnight cultures of cells freshly transformed with plasmid(s) were diluted 1:40 into fresh medium supplemented with appropriate antibiotic(s) as indicated below. For export assays of MdoD linker variants, cultures were grown in LB containing ampicillin (100 μg/ml) for 45 minutes at 37°C, IPTG was added to a final concentration of 1 mM and cultures were grown for a further 45 minutes. For MdoD signal peptide deletion experiments, cultures were grown in LB with 1 mM IPTG for 45 min at 37°C, L arabinose was added to a final concentration of 0.2% L-arabinose and cultures were grown for a further 45 minutes. Cells were harvested by centrifugation (3250*g*, 4°C, 10 min) and resuspended in 10 mM Tris-HCl, 150 mM NaCl, pH 7.3 with cell densities normalised according to A_600nm_. Equal volumes of the resuspended cells were collected by centrifugation (16,000*g*, 4°C, 1 min) and resuspended in 400 μl SET buffer (17% sucrose (w/v), 3 mM EDTA, 10 mM Tris.HCl pH 7.3). 133 μl of lysozyme (3 mg/ml) and 400 μl of ice-cold water were added and samples incubated at 37°C for 20 min. Spheroplasts were separated from the periplasmic contents by centrifugation (16,000*g*, 4°C, 1 min). Spheroplasts were washed in 10 mM Tris.HCl, 150 mM NaCl, pH 7.3 and collected by centrifugation (16,000*g*, 4°C, 1 min). Samples were analysed by immunoblotting for MdoD in the linker deletion experiments using monoclonal anti-Myc antibody. Samples were analysed by immunoblotting for MdoD in the signal peptide deletion experiments using monoclonal anti-FLAG antibody (full length C-terminally FLAG tagged MdoD (MdoD_wild type_)) or using monoclonal anti-Myc antibody (MdoD_33-351_ (MdoD_Δsignal_), C-terminally Myc tagged). Polyclonal GroEL antibody was used to detect GroEL as a cytoplasmic control.

### Affinity chromatography co-purification assays

Purified full length His_6_-SUMO-CueO_myc_ and His_6_-SUMO-MdoD_myc_ (purified as described above) were incubated with *Saccharomyces cerevisiae* Ubiquitin-like-specific protease 1 (Ulp1) at 4°C for 12 hours and injected onto a Superose 6 Increase 10/300 GL size exclusion column (cytiva) pre-equilibrated in buffer C (20 mM HEPES pH 7.4, 150 mM NaCl, 0.02% GDN) to separate His-tagged SUMO from CueO or MdoD. Fractions corresponding to CueO or MdoD were collected and stored on ice and used the same day. Membranes containing overexpressed His tagged *E. coli* TatBC wild type or its variants were prepared and solubilised in GDN as described above. Insoluble material was removed by centrifugation (100,000*g*, 30 min) and clarified solubilised membranes were incubated with 200 ul of pre-washed cOmplete His-tag purification resin (Roche, 50% w/v slurry). The resin was washed with 15 CV of buffer (50 mM sodium phosphate pH 8.0, 150 mM NaCl, 2 mM imidazole, 1 mM EDTA, 0.02% (w/v) GDN).

Purified CueO was incubated with resin bound *E. coli* TatBC for 2 hours. The resin was washed with 30 CV of buffer (50 mM sodium phosphate pH 8.0, 150 mM NaCl, 2 mM imidazole, 1 mM EDTA, 0.02% (w/v) GDN). Proteins were eluted from the resin with buffer (50 mM sodium phosphate pH 8.0, 150 mM NaCl, 200 mM imidazole, 1 mM EDTA, 0.02% (w/v) GDN). Proteins were separated by SDS–PAGE and detected by immunoblotting using polyclonal antibodies against TatB, TatC and CueO.

*N. salsuginis* TatABC, TatBC and TatAC complexes for use in affinity chromatography co-purification assays were generated as described below. *N. salsuginis* TatABC was overexpressed in *E. coli* strain BL21 DE3 carrying both a pRSFDuet-1 vector containing *tatC*-strep (mcs1) and *tatB*-flag (mcs2) and a pACT7 vector containing *tatA*-(His)_6_. *N. salsuginis* TatBC was overexpressed in *E. coli* strain BL21 DE3 carrying the pRSFDuet-1 vector containing *tatC*-strep (mcs1) and *tatB*-flag (mcs2). *N. salsuginis* TatAC was overexpressed in *E. coli* strain BL21 DE3 carrying both a pRSFDuet-1 vector containing only *tatC*-strep (mcs1) and a pACT7 vector containing *tatA*-(His)_6_. Cells were grown in TB media containing Kanamycin (50 μg/ml) and chloramphenicol (20 μg/ml) at 37°C with shaking to A_600nm_ = 4.0 after which cultures were supplemented with 0.1 mM IPTG and incubated for an additional 12 h at 25°C with shaking. Cells were harvested by centrifugation (5,000*g,* 10 min). Membranes were prepared and solubilised in GDN as described above. Insoluble material was removed by centrifugation (100,000*g*, 30 min) and clarified solubilised membranes were incubated with 200 ul of pre-washed streptactin XT superflow resin (50% w/v slurry, IBA Lifesciences). The resin was washed with 15 CV of buffer (50 mM sodium phosphate pH 8.0, 150 mM NaCl, 1 mM EDTA, 0.02% (w/v) GDN). Purified CueO or MdoD were incubated with resin bound *N. salsuginis* TatABC, TatBC or TatAC for 2 hours. The resin was washed with 30 CV of buffer (50 mM sodium phosphate pH 8.0, 150 mM NaCl, 1 mM EDTA, 0.02% (w/v) GDN). Proteins were eluted from the resin with buffer (50 mM sodium phosphate pH 8.0, 150 mM NaCl, 50 mM D-biotin, 1 mM EDTA, 0.02% (w/v) GDN). Proteins were separated by SDS–PAGE and detected by immunoblotting using antibodies against His-tag (TatA), FLAG-tag (TatB), strep tag (TatC) or Myc tag (MdoD or CueO).

### Gel quantification to determine TatB:TatC ratios

Purified TatABC and TatBC were loaded onto 4-20% SDS-PAGE gels at the following dilutions: 1, 0.75, 0.5 and 0.25. Gels were run at 180 V for 30-40 min and stained with Coomassie brilliant blue. Gels were imaged with a ChemiDoc MP imaging system (Bio-Rad) and gel bands were quantified using ImageJ. Quantification was performed with samples from three independent biological repeats.

### Coarse-grained molecular dynamics simulations

Membrane thinning was investigated using coarse-grained (CG) simulations. The E. coli TatBC complex cryo-EM structure was used as an input. The input protein was aligned according to the plane of the membrane with MEMEMBED(*85*) and converted to a CG representation using the Martini 3 force field(*86*). An elastic network with a force constant of 500 kJ mol-1 nm-2 was used to restrain the secondary structure of the protein. A POPE:POPG bilayer at 1:4 molar ratio was built around the protein using the insane protocol(*87*). The system was solvated with Martini 3 waters and NaCl was added in a concentration of 150 mM to neutralize the system. The system was energy minimized using the steepest descents method, followed by 10 ns equilibration in the NVT ensemble, then by 10 ns in the NPT ensemble, before 3 x 5μ*s* production simulations. All production simulations sampled isothermic-isobaric ensembles at 310 K using the V-rescale thermostat (τ_*T*_= 1.0)(*88*), the C-rescale barostat for semi-isotropic pressure coupling at 1.0 bar (τ_P_= 12.0)(*89*) and a time-step of 20 fs. The Reaction-Field method was used to model long range electrostatic interactions. Bond lengths were constrained to their equilibrium values.

### Atomistic simulations

The cryo-EM structures of the TatBC complex, in the presence and absence of the MdoD signal peptide, were used as starting points for atomistic simulations. To model the APH of TatA, which is absent in the cryo-EM structure, the TatBC complex was used. Similarly, the TatBC– MdoD structure served as the basis for modeling the TatAC–MdoD complex. Atomistic simulations for all four system were prepared using the MemProtMD(*90*) pipeline. After 1μ*s* the CG systems were converted back to atomistic details using CG2AT(*91*). The systems were further equilibrated for 1ns maintaining the structure of the protein restrained. Three repeats of unrestrained 500 ns MD simulations were performed for each system. All simulations were performed in the isothermal-isobaric ensemble at 310 K and 1 bar using a time-step of 2 fs and the CHARMM36 force field(*92*). Pressure was maintained at 1 bar using the C-rescale barostat (τ_P_ = 1.0). Temperature was controlled using the velocity rescale thermostat (τ_*T*_ = 0.1), with the solvent, lipids and protein coupled to an external bath. Electrostatics was described using the Particle-mesh Ewald method(*93*), with a cutoff of 1.2 nm, and the Van der Waals interactions were shifted between 1 and 1.2 nm. All MD simulations were performed using GROMACS 2023.4(*94, 95*).

## Author contributions

**Justin C. Deme:** Conceptualization, Investigation, Writing-Original Draft, Visualisation, Methodology **Owain J. Bryant:** Conceptualization, Investigation, Writing-Original Draft, Visualisation, Methodology **Mariana R. B. Batista:** Conceptualization, Investigation, Writing, Visualisation, Methodology related to MD **Phillip J. Stansfeld**: Conceptualization, Writing, Methodology, Supervision, Funding Acquisition related to MD **Ben C. Berks**: Conceptualization, Writing-Original Draft, Methodology, Supervision, Funding Acquisition **Susan M. Lea**: Conceptualization, Writing-Original Draft, Visualisation, Methodology, Supervision, Funding Acquisition

## Data availability

Cryo-EM reconstructions and atomic models for *E. coli* TatBC (EMD-47343; PDB 9DZZ), *N. salsuginis* TatBC (EMD-47346; PDB 9E02), *M. xanthus* TatBC (EMD-47347; PDB 9E03), *E. coli* TatBC-MdoD (EMD-47345; PDB 9E01), *E. coli* MdoD (EMD-47352; PDB 9E08), *N. salsuginis* TatBC-CueO (EMD-47348; PDB 9E04), *N. salsuginis* TatBC in nanodisc (EMD-47350; PDB 9E06), *N. salsuginis* TatAC (EMD-4735; PDB 9E07) have been deposited in the Electron Microscopy Data Bank and Protein Data Bank, respectively, with the appropriate accession codes listed in parenthesis beside each entry. The atomic model of *M. xanthus* TatC periplasmic domain solved by X-ray crystallography was deposited in the Protein Data Bank with accession code 9E0S.

## Acknowledgments

We acknowledge the use of the Central Oxford Structural Microscopy and Imaging Centre (COSMIC) and the CCR Center for Structural Biology CryoEM core. This work was supported by Wellcome Trust Investigator Award 107929/Z/15/Z (B.C.B) and Medical Research Council grant MR/L000776/1 (B.C.B and S.M.L.) and MR/S021264/1 (S.M.L.). This research was supported in part by the Intramural Research Program of the NIH (S.M.L.). PJS’s lab was funded by Wellcome (208361/Z/17/Z), MRC, BBSRC, EPSRC, NIH, JPIAMR and the Howard Dalton Centre. We acknowledge the use of time on ARCHER2 granted via the UK High-End Computing Consortium for Biomolecular Simulation, HECBioSim (http://www.hecbiosim.ac.uk), supported by EPSRC grant no. EP/R029407/1. We acknowledge the use of time on Sulis funded by EPSRC grant no. EP/T022108/1. We acknowledge the use of the SCRTP at Warwick computing infrastructure.

## List of Supplementary Materials

## Supplementary Materials For

**Fig S1.**
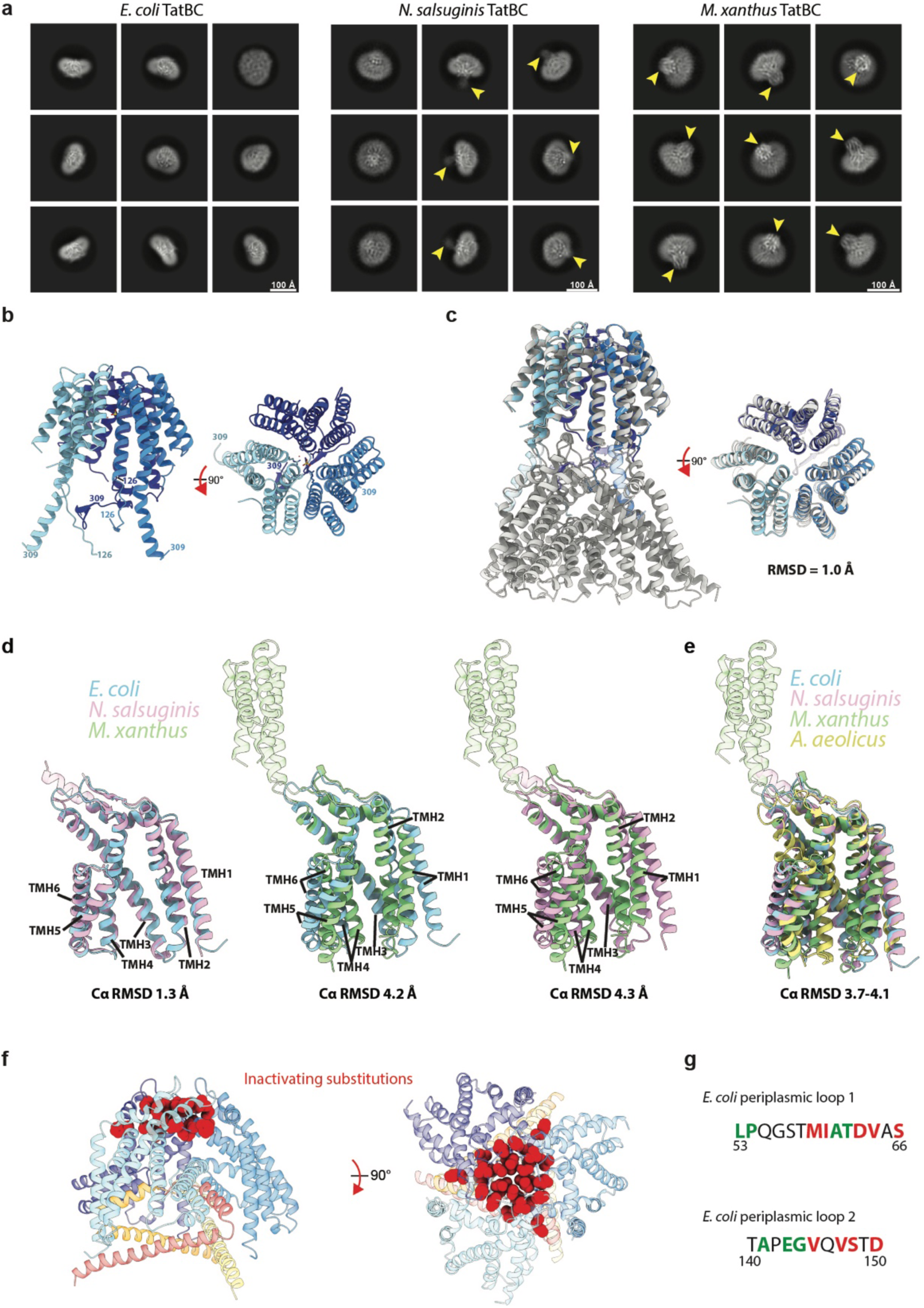
Architectures of TatBC complexes. **a**, Cryo-EM 2D class averages of TatBC complexes purified in GDN from *E. coli* (left), *N. salsuginis* (middle), and *M. xanthus* (right). The yellow arrowheads indicate the location of the periplasmic domains in the *N. salsuginis* and *M. xanthus* complexes. **b,** Cartoon representation of the structure of the isolated periplasmic domain of *M. xanthus* determined by X-ray crystallography with each subunit coloured in different shades of blue. A sulphate atom bound on the three-fold axis and coordinating side chains and waters shown in stick representation coloured by chemical composition **c,** Structural alignment of the *M. xanthus* periplasmic domain solved by X-ray crystallography (various shades of blue) and the TatBC complex solved by cryo-EM (grey). Residues 135-289 were used for alignment. Those parts of the crystal structure not used in the alignment are shown transparent. **d**, Pairwise structural alignments of the TatC models presented in this study. **e**, Structural alignment of the TatC models presented in this study against the crystal structure of *A. aeolicus* TatC (yellow; PDB 4B4A). Periplasmic residues that did not contribute to the alignment are shown with increased transparency. **f,** Model of the *E. coli* TatBC complex viewed from the plane of the membrane (left) or the periplasm (right). Residues (red space filling representation) within the periplasmic loops of TatC where mutations have previously been shown to be important for core complex function or assembly^1,2^. **g,** Positions within the TatC periplasmic loops coloured according to whether previously reportedsubstitutions^1,3,4^ inactivate (red) or have no effect (green) on Tat transport. Positions not tested are coloured black.

**Fig S2.**
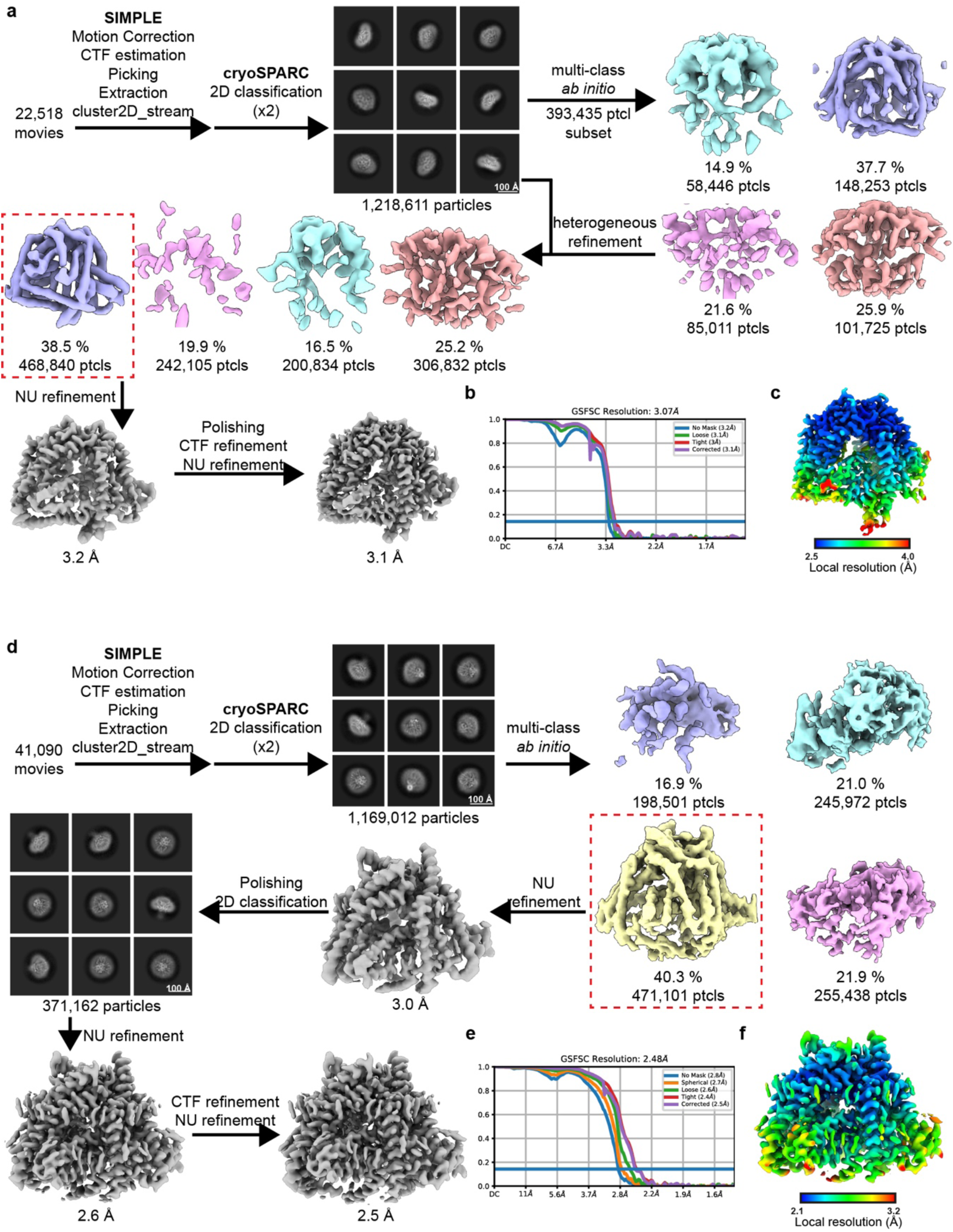
Cryo-EM workflows for *E. coli* and *N. salsuginis* TatBC complexes. *E. coli*. **(a-c)** and *N. salsuginis* **(d-f)** cryoEM workflows. **a,d,** Image processing workflow. **b,e,** Gold-standard Fourier Shell Correlation (FSC) curves used for global resolution estimation. **c,f,** Local resolution estimate of the volume.

**Fig S3.**
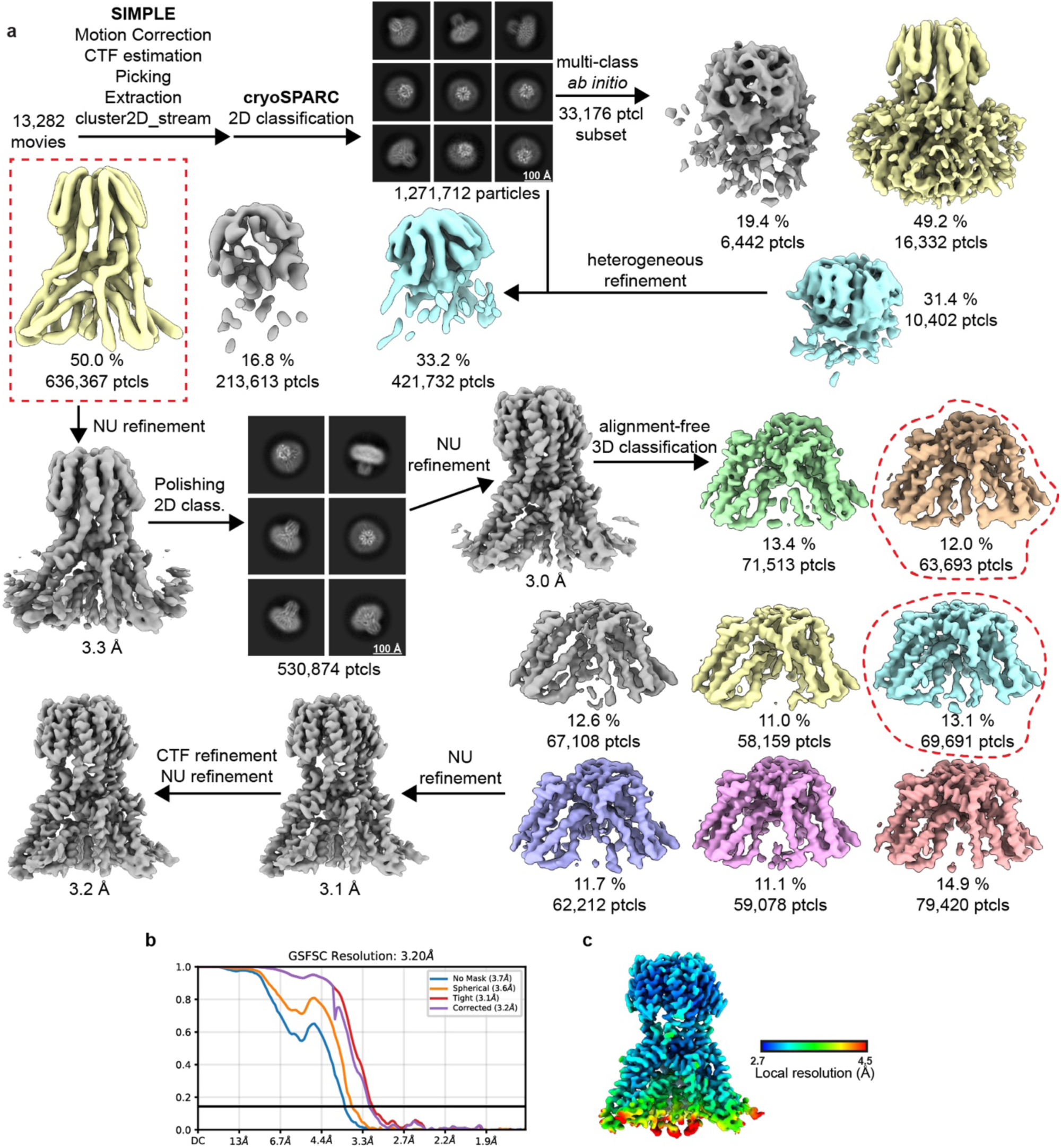
Cryo-EM workflow for the *M. xanthus* TatBC complex. **a**, Image processing workflow. **b,** Gold-standard Fourier Shell Correlation (FSC) curves used for global resolution estimation. **c,** Local resolution estimate of the volume.

**Fig S4.**
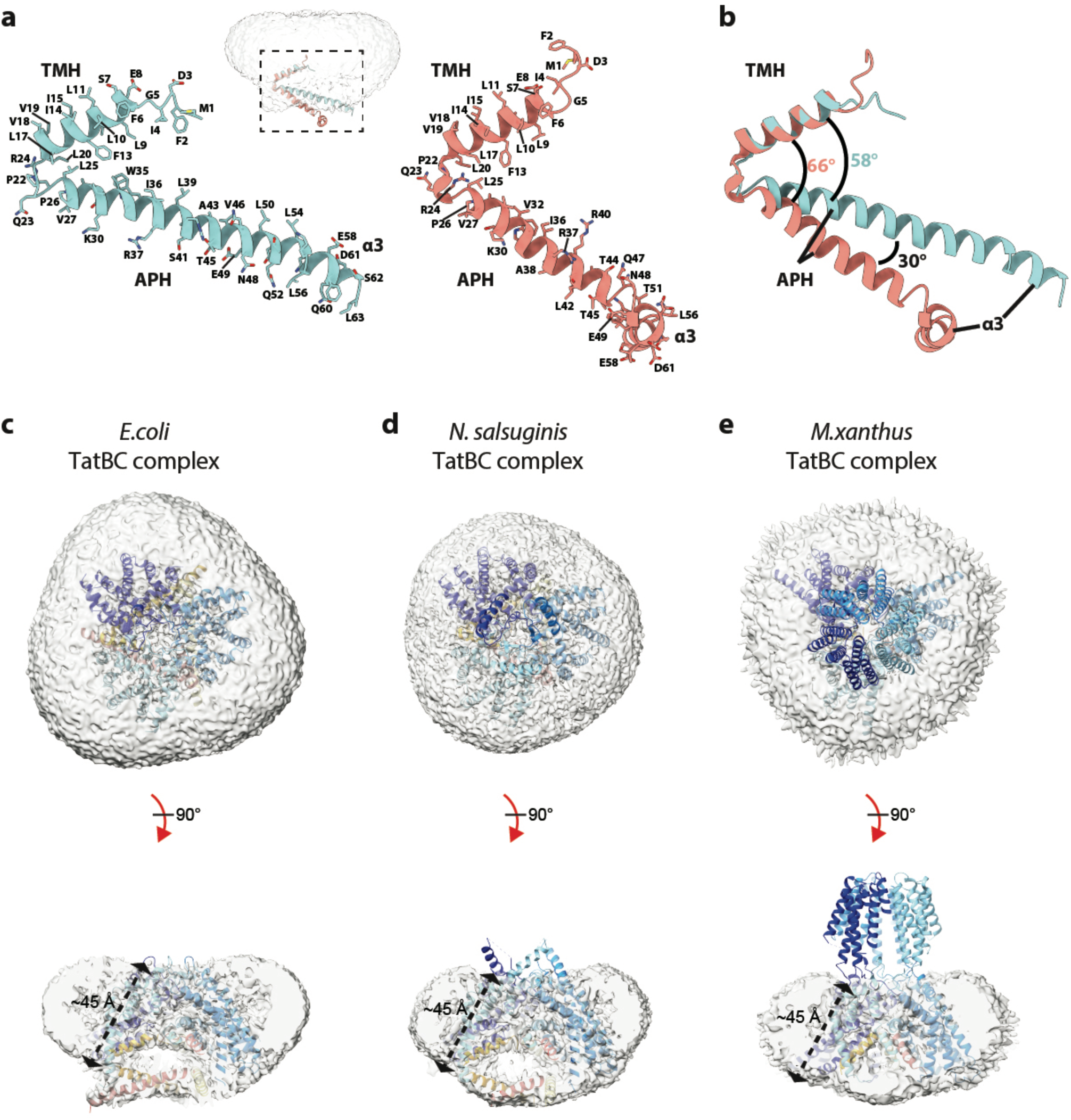
TMH tilting leads to thinning of the detergent micelle around the TatBC complex. **a**,**b,** The arrangement of the TatB helices observed in the TatBC complex (sky blue) differs from that in an earlier NMR structure of isolated TatB (salmon, PDB 2MI2). The structures are shown separately (a) or superimposed (b, aligned on TMH residues 6-21, Cα RMSD ∼ 0.5 Å). The inset in (a) shows the NMR structure superimposed on the position of TatB in the TatBC complex. Alignments and RMSD were calculated in ChimeraX^5^. **c-e,** Tilting of the TatC TMHs thins the hydrophobic layer around the TatBC complex as seen in the detergent/phospholipid micelles around the purified complexes. Models of the *E. coli* (**c**), *N. salsuginis* (**d**) and *M. xanthus* (**e**) TatBC complexes superposed with low contoured cryo-EM volumes to display the detergent/phospholipid micelle (grey). TatB density is depicted in red, orange and yellow, TatC density in various shades of blue.

**Fig. S5.**
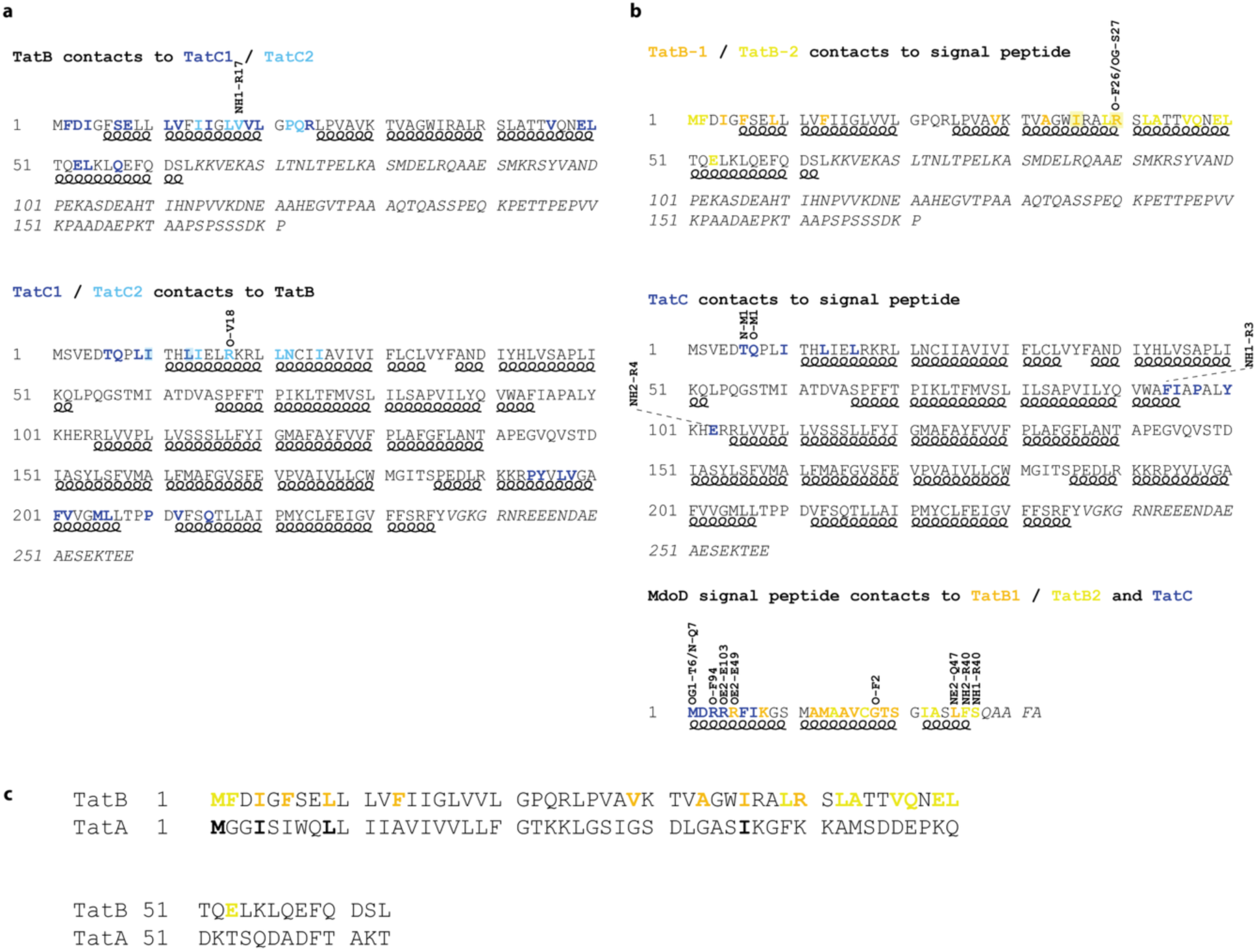
Mapping *E. coli* core complex interactions onto the protein sequences. Residues involved in helices are denoted by helix beneath the sequence. Residues not seen in the structures are in italics. Colour indicates which chain is involved in the contact. **a**, Contacts between *E. coli* TatB and TatC. Coloured residues indicate which copy of TatC is contacted. Atoms in the other chain to which the coloured residue forms a hydrogen-bond are shown vertically. **b**, Contacts to the MdoD signal peptide. Coloured residues are o denote which copy of TatB is involved and the signal peptide residues are coloured to denote the Tat chain contacted. **c**, TatA and TatB are aligned from the N-terminus with the ordered portion of TatB shown. Residues coloured in TatB are involved in contacts with signal peptide. Of these 15 TatB residues only 4 are identical in TatA (indicated in bold).

**Fig. S6.**
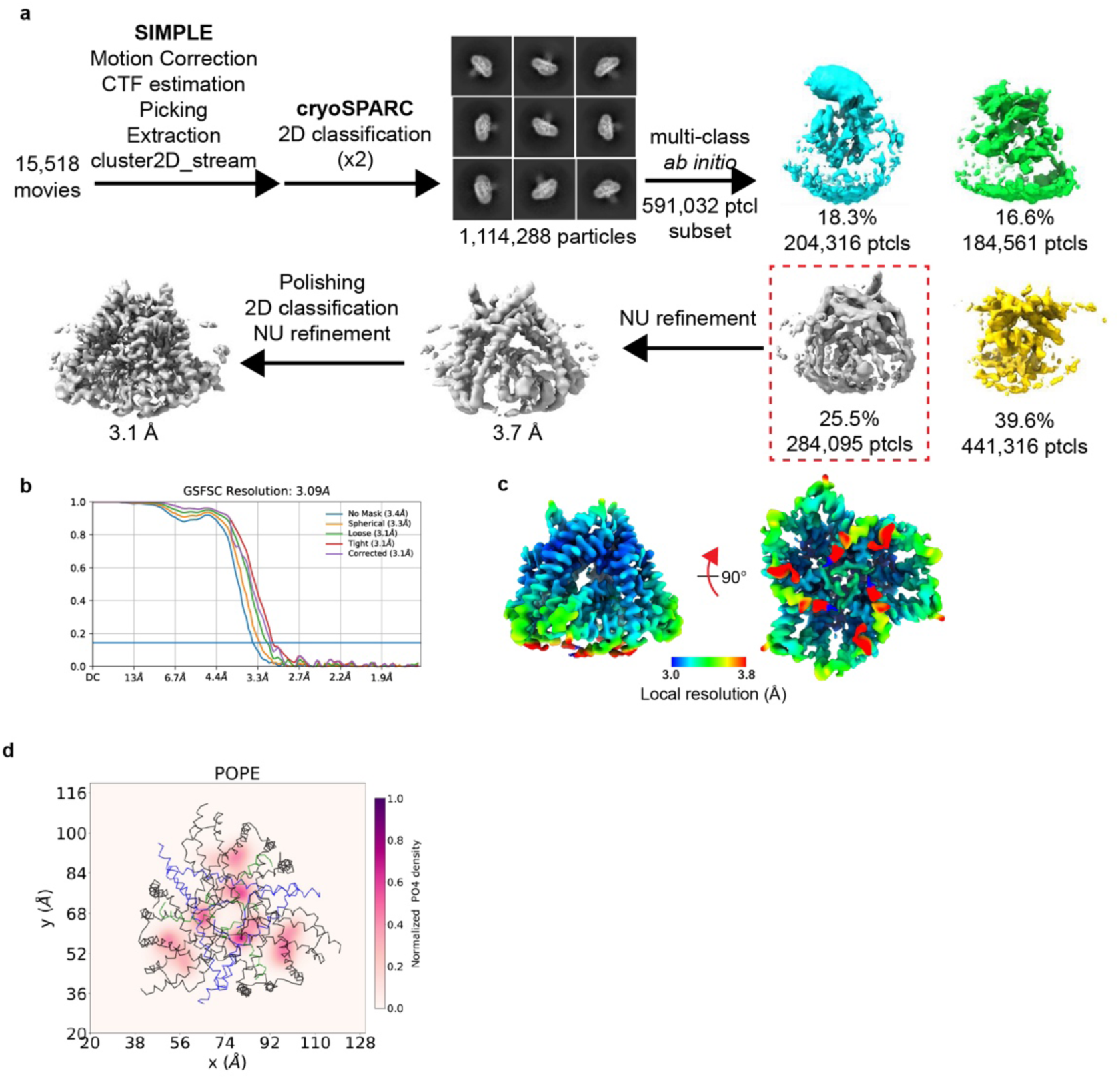
Cryo-EM workflow for the *N. salsuginis* TatBC complex in a nanodisc and the distribution of lipids in an MD simulation of *E. coli* TatBC. **a-c**, CryoEM workflow for the *N. salsuginis* TatBC complex in a nanodisc. **a,** Image processing workflow. **b,** Local resolution estimate of the volume. **c,** Gold-standard Fourier Shell Correlation (FSC) curves used for global resolution estimation. **d,** Density plot of the lipids located within the TatBC complex. The lipids remain stably associated within the complex and do not diffuse into the bulk membrane.

**Fig S7.**
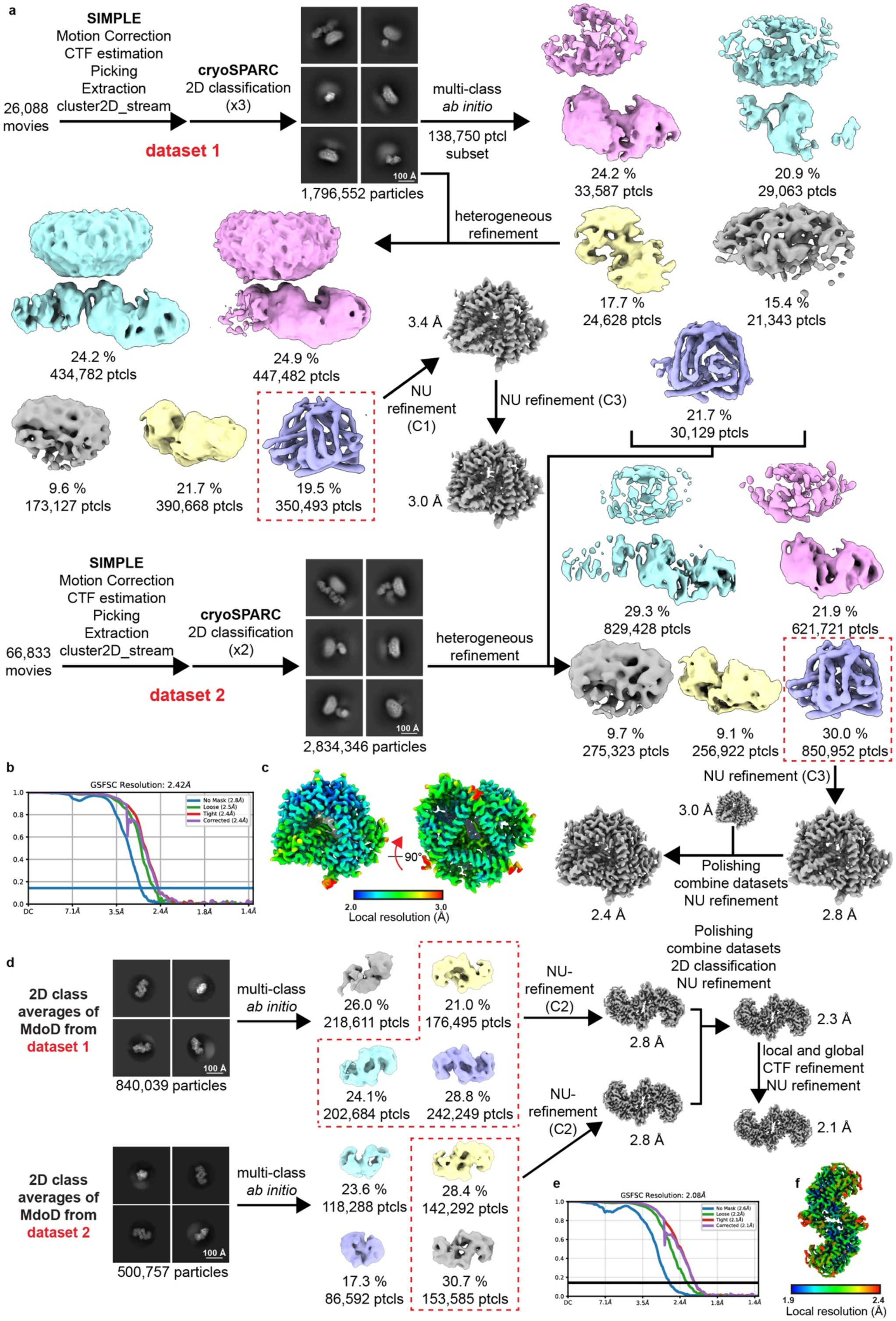
Cryo-EM workflow for the *E.coli* TatBC–MdoD complex and MdoD dimer. *E. coli* TatBC–MdoD complex. **(a-c)** and MdoD dimer **(d-f)** cryoEM workflows. **a,d,** Image processing workflow. **b,e,** Gold-standard Fourier Shell Correlation (FSC) curves used for global resolution estimation. **c,f,** Local resolution estimate of the volume.

**Fig. S8.**
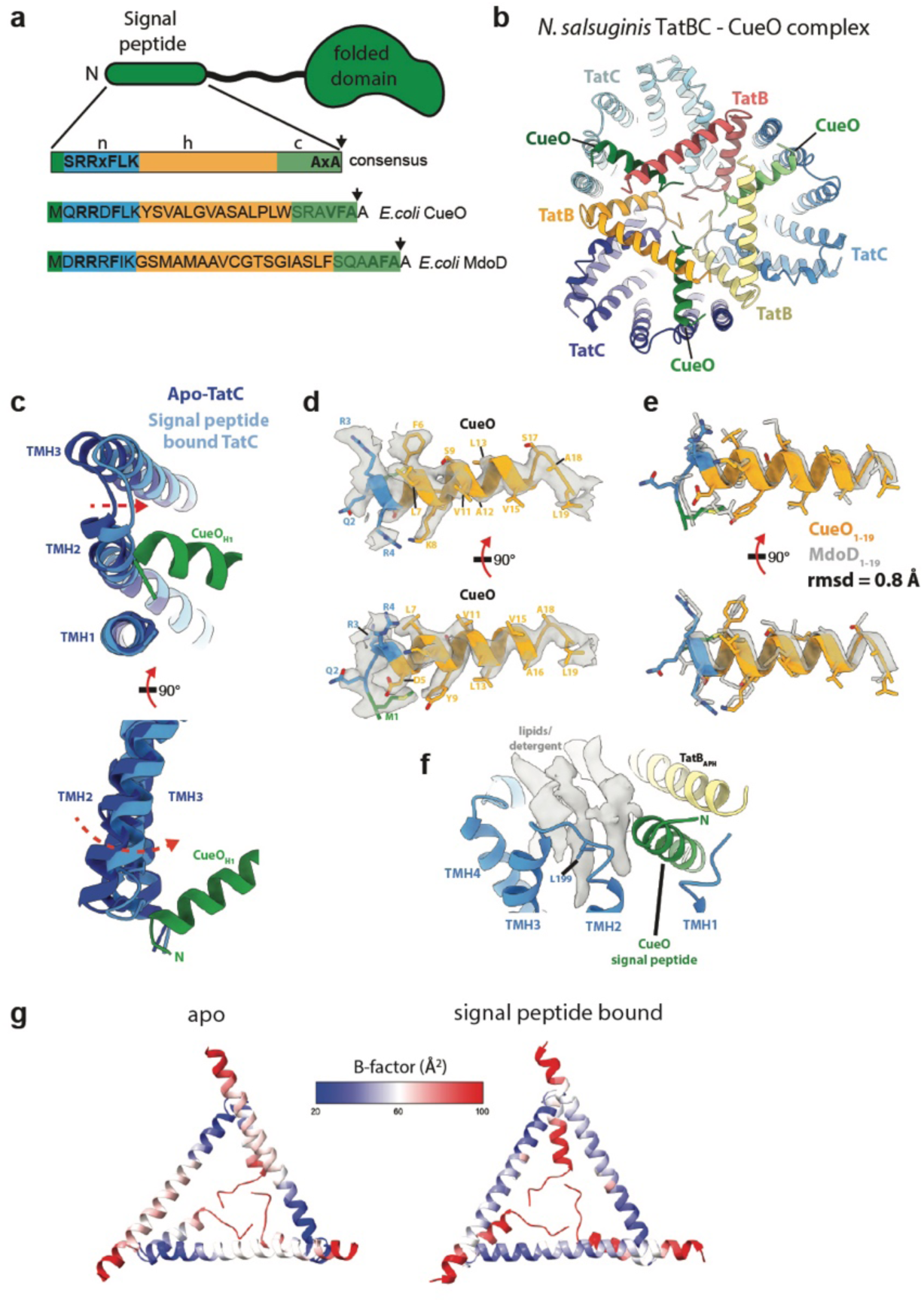
Signal peptide recognition by the TatBC complex. **a**, The features of a Tat signal peptide shown schematically (top) and identified within the sequences of the signal peptides of the *E. coli* proteins CueO and MdoD (below). The site of cleavage by signal peptidase is indicated by a black arrow. **b-f**, Structural analysis of the *N. salsuginis* TatBC-CueO signal peptide complex. **b,** Model of the complex viewed from the cytoplasm. **c,** Structural alignment of the signal peptide-bound (TatC light blue and signal peptide green) and apo (TatC dark blue) TatBC complexes (arrow shows movements on signal peptide binding). The selected portion of the complex is viewed from the cytoplasm (top) or the membrane plane (bottom). **d,** Model for the CueO signal peptide shown in the cryo-EM volume (transparent grey surface) with the peptide coloured as in (a). **e,** Structural alignment of the CueO signal peptide from the *N. salsuginis* TatBC-CueO signal peptide complex against the MdoD signal peptide from the *E. coli* TatBC-MdoD complex. Alignments and RMSD were calculated using the MatchMaker command in ChimeraX^5^. **f**, Lipids are packed between TatC and the signal peptide. *N. salsuginis*Leu199 (in sticks representation) is the equivalent of *E. coli* Leu99 (Fig. 3h). **g**, The ordering of the TatB APH increases on signal peptide binding. Cartoon representation of TatB coloured by B factors for the apo (left) and MdoD-bound (right) *E. coli* TatBC complexes.

**Fig. S9.**
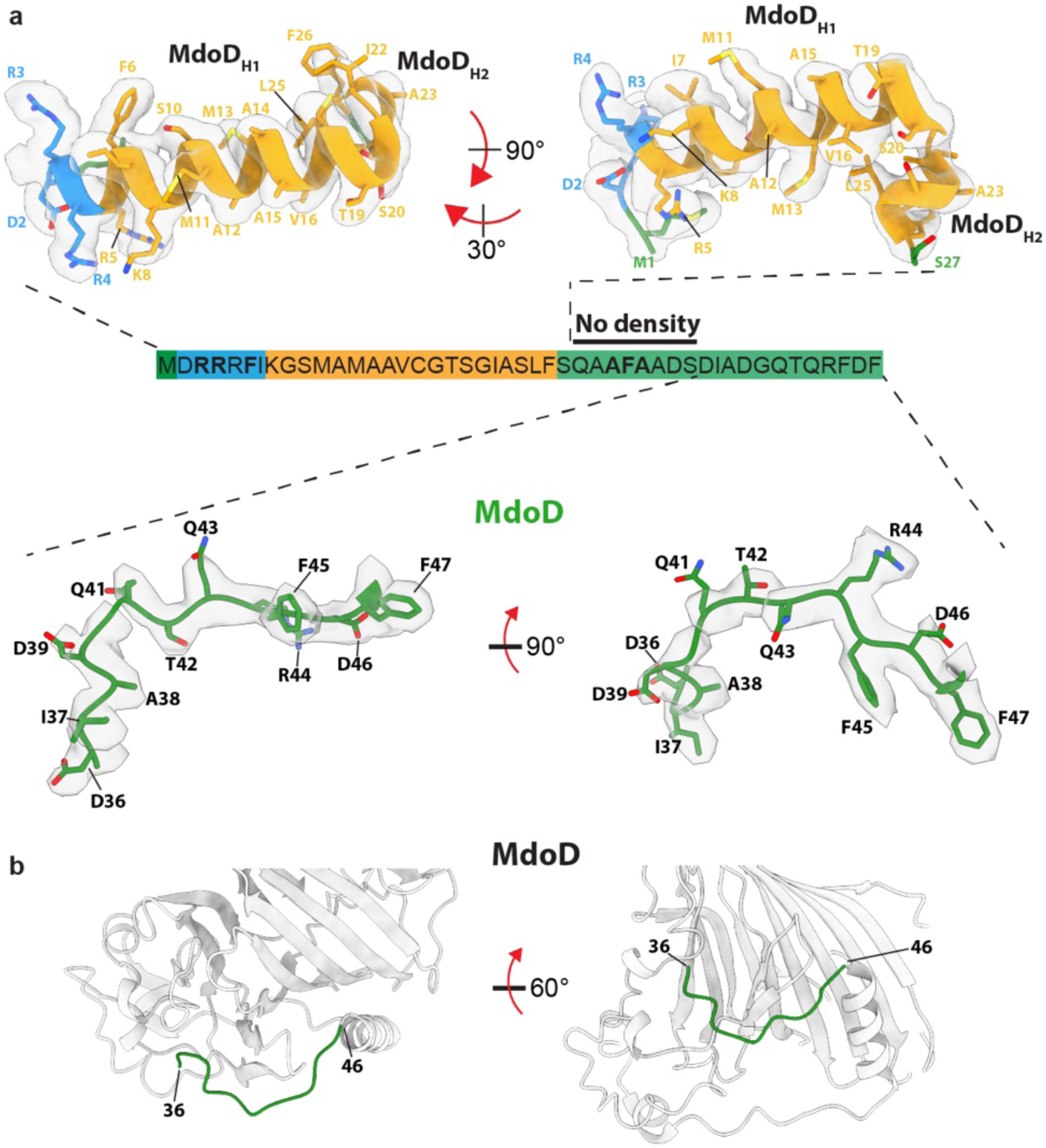
The disordered linker between the signal peptide and folded domain of MdoD extends from residue 28 to 35. **a**, The signal peptide (above) and first ten residues of the folded domain (below) are shown in their respective cryo-EM volumes (semi-transparent, grey). The region between these residues is not present in either the membrane domain-focused or folded substrate domain-focused cryo-EM volumes and is presumed disordered. (Centre) The amino acid sequence corresponding to MdoD residues 1 – 47. Residues where density is not visible in the TatBC-signal peptide volume or MdoD folded domain volume are highlighted. The protein models are coloured according to the schematic. **b**, The location of MdoD residues 36-46 (green) which pack against the rest of the folded domain of MdoD (light grey).

**Fig. S10.**
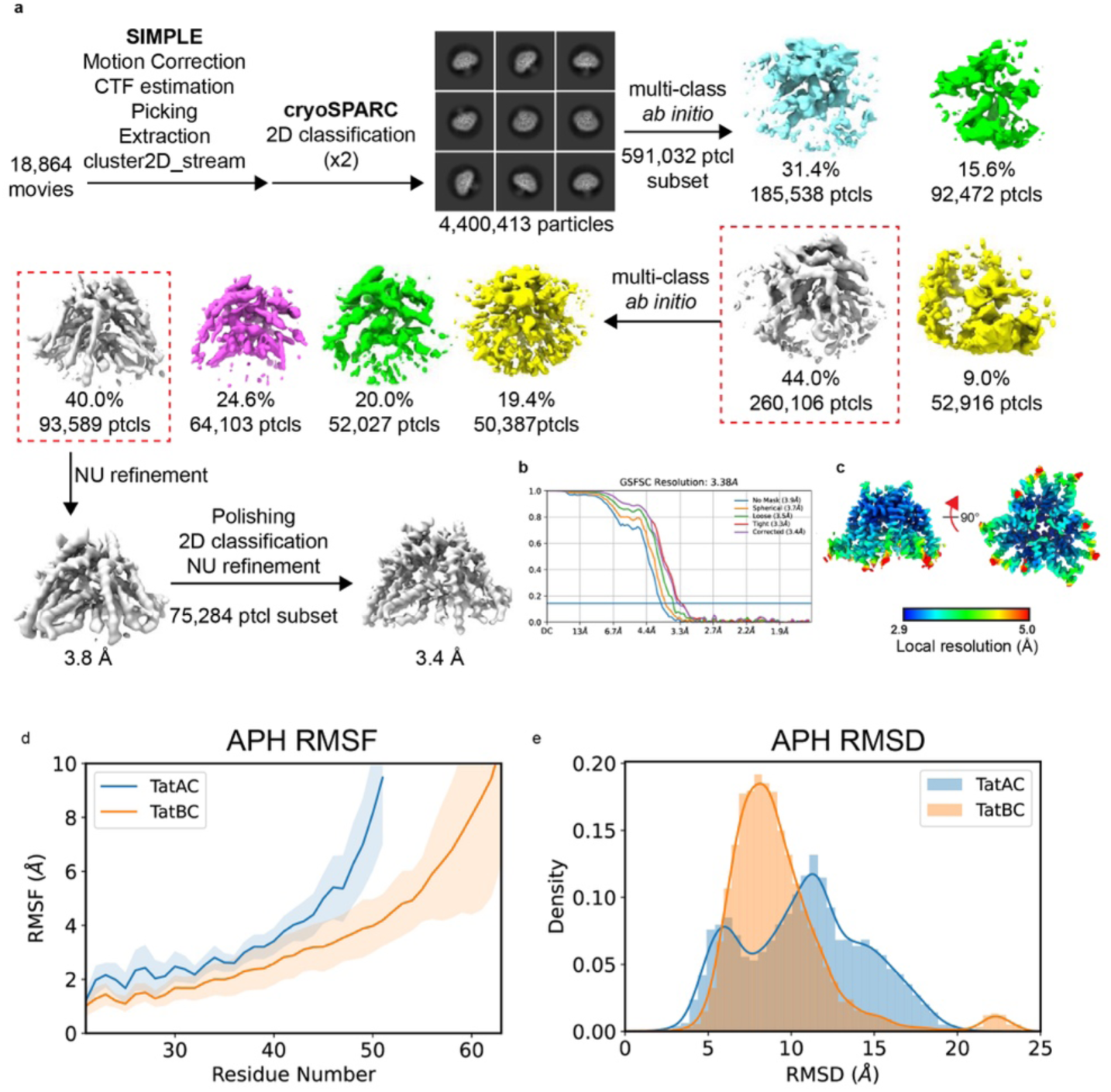
Cryo-EM workflow for the *N. salsuginis* TatAC complex and simulations of TatA complexes. **a-c** CryoEM workflow for the *N. salsuginis* TatAC complex. **a,** Image processing workflow. **b,** Gold-standard Fourier Shell Correlation (FSC) curves used for global resolution estimation. **c,** Local resolution estimate of the volume. **d,** Root Mean Square Fluctuation (RMSF) of the amphipathic helix of TatA and TatB, indicating differences in flexibility along the helix residu. **e,** TatA and TatB APH RMSD distributions. The RMSD and RMSF were calculated using TatA/TatB TMH as reference to align the MD frames.

**Fig. S11.**
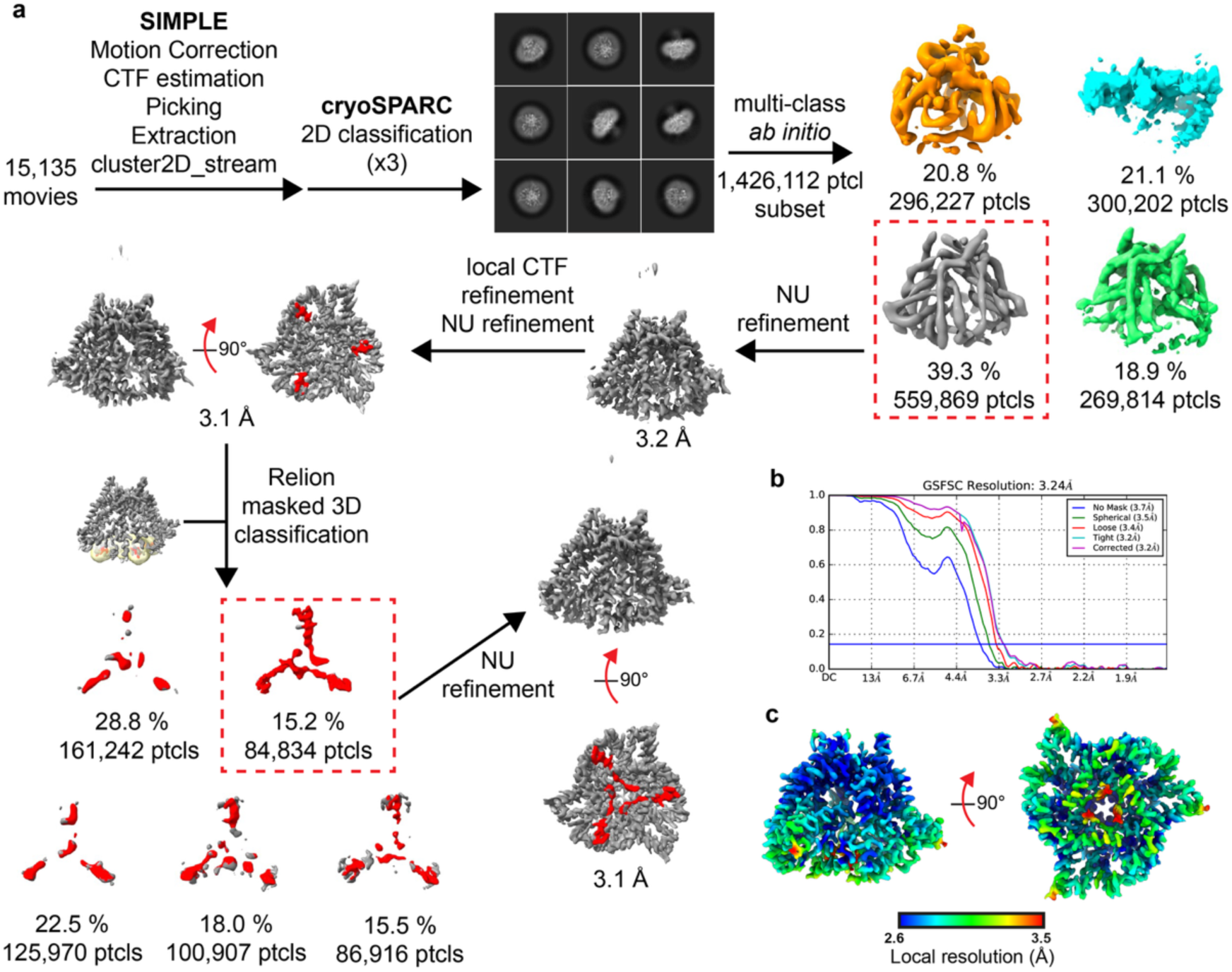
Cryo-EM workflow for *N. salsuginis* TatBC-CueO signal peptide complex. **a**, Image processing workflow. **b**, Gold-standard Fourier Shell Correlation (FSC) curves used for global resolution estimation. **c**, Local resolution estimation of the volume

**Fig. S12.**
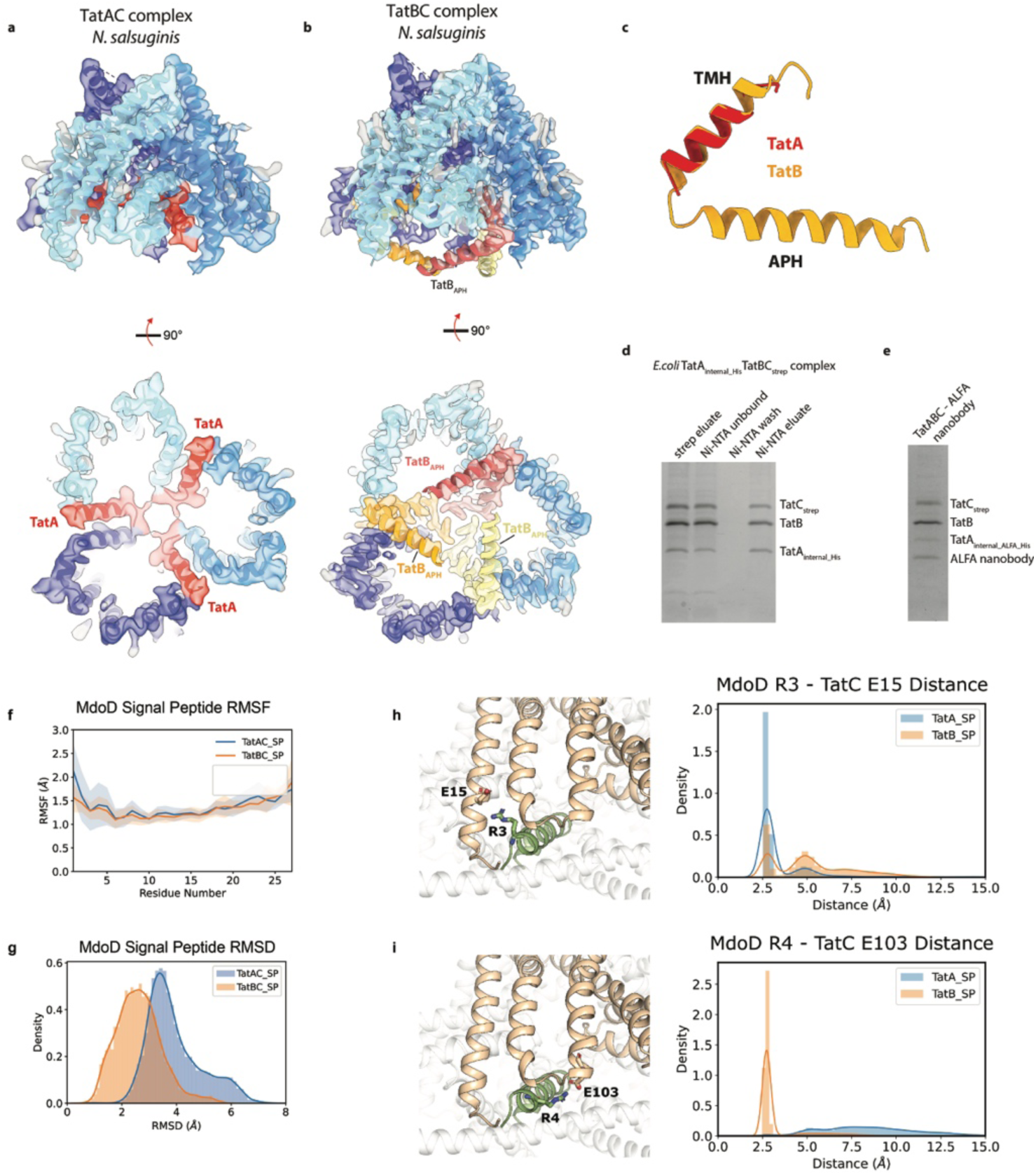
Incorporation of TatA into the Tat core complex complex and simulations of signal peptide contacts. **a, b**, Models of *the N. salsuginis* TatAC (a) and TatBC (b) complexes shown as cartoon representations (TatC-blue, TatA-red, TatB-yellow/orange/red) in their respective cryoEM volumes (semi-transparent) viewed from the plane of the membrane (top) or the cytoplasm (bottom) Structures behind the view plane in the view from the cytoplasm, have been removed to simplify the view. **c**, Superposition of the structurally defined portions of the TatA (red) and TatB (gold) subunits in the *N. salsuginis* TatAC and TatBC structures following alignment on the TatC subunits (Cα RMSD ∼ 0.6 Å). Alignments and RMSD were calculated using the MatchMaker command in ChimeraX^5^. **d,** Isolation of an *E. coli* TatA_His_TatBC_strep_ complex solubilized in GDN and purified successively by strep tag affinity chromatography and Ni-NTA affinity chromatography. Th indicated samples are analysed by SDS-PAGE and stained with Coomassie Blue. **e,** Nanobody labelling of the TatA subunit within the TatABC complex. An ALFA nanobody was mixed with purified *E. coli* TatA_ALFA,His_TatBC_strep_ complex and subject to separation by size exclusion chromatography. Coomassie Blue stained SDS-PAGE gel of the fraction from the size exclusion column used for cryoEM. **f,** RMSF and **g**, RMSD distributions of the signal peptide when bound to TatAC and TatBC complexes. RMSF and RMSD were calculated using TatC as reference for aligning the MD simulation frames. **h**, MdoD R3 – TatC E15 and **i**, MdoD R4 – TatC E103 distance distributions. Distances were defined as the minimum distance between the hydrogen bond donor/acceptor atoms. global resolution estimation. **c**, Local resolution estimation of the volume

**Fig. S13.**
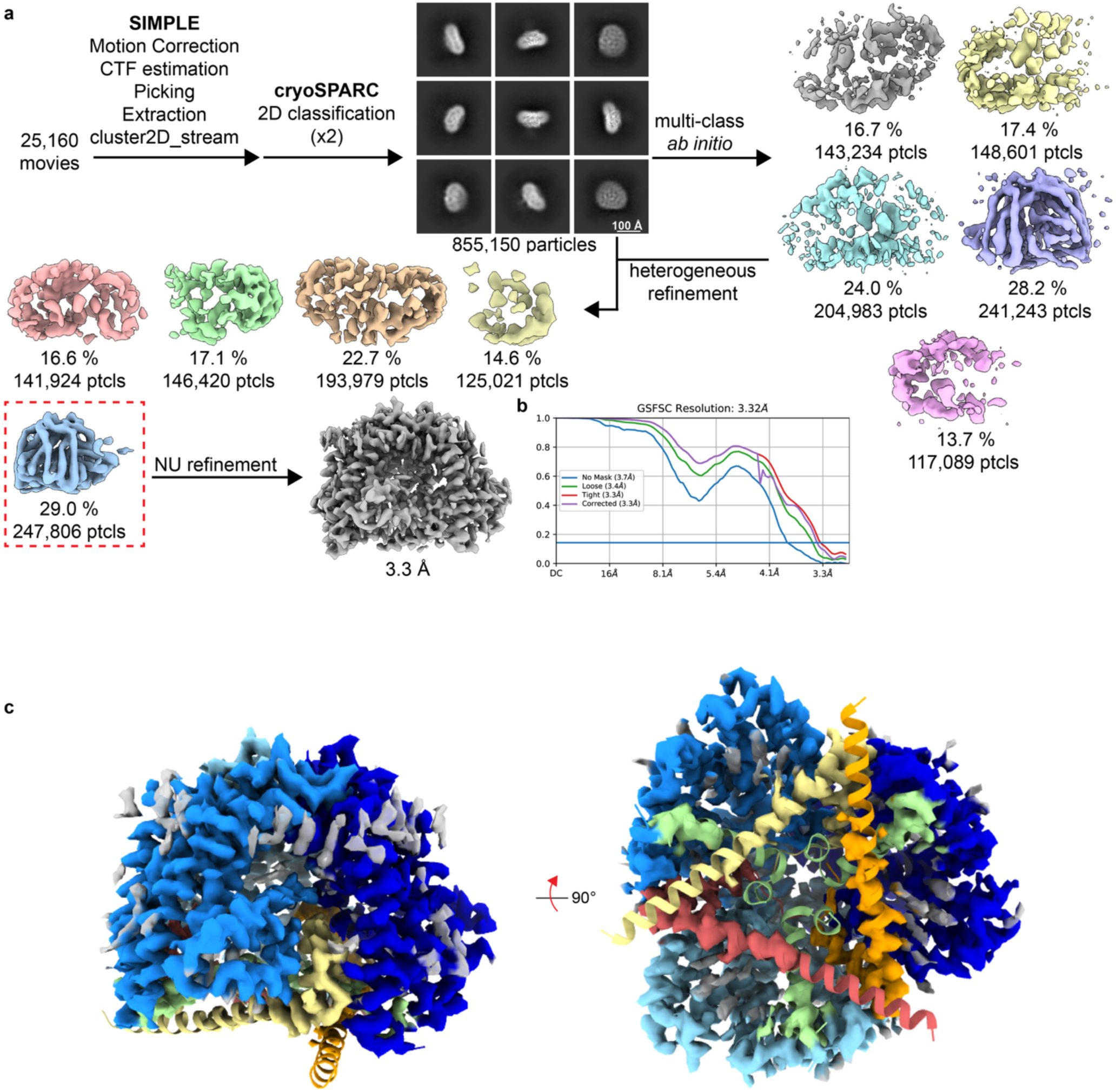
Asymmetric C1 reconstruction of an *E. coli* TatABC-substrate complex. An unsymmetrized C1 reconstruction. The TatABC complex was purified using an internal his-tag at position 26 of TatA. to which CueO substrate was added prior to grid preparation. **a**, Image processing workflow. **b**, Gold-standard Fourier Shell Correlation (FSC) curves used for global resolution estimation. **c**, The C1 volume is not appreciably asymmetric at the resolution of the complex (3.3A). The volume is essentially indistinguishable from the TatBC-MdoD complex (coordinates shown as docked cartoon representation) except that the amphipathic helices of TatB and the substrate peptide are substantially less well ordered in the TatABC-CueO volume implying lower occupancy and/or higher mobility of these regions.

**Fig. S14.**
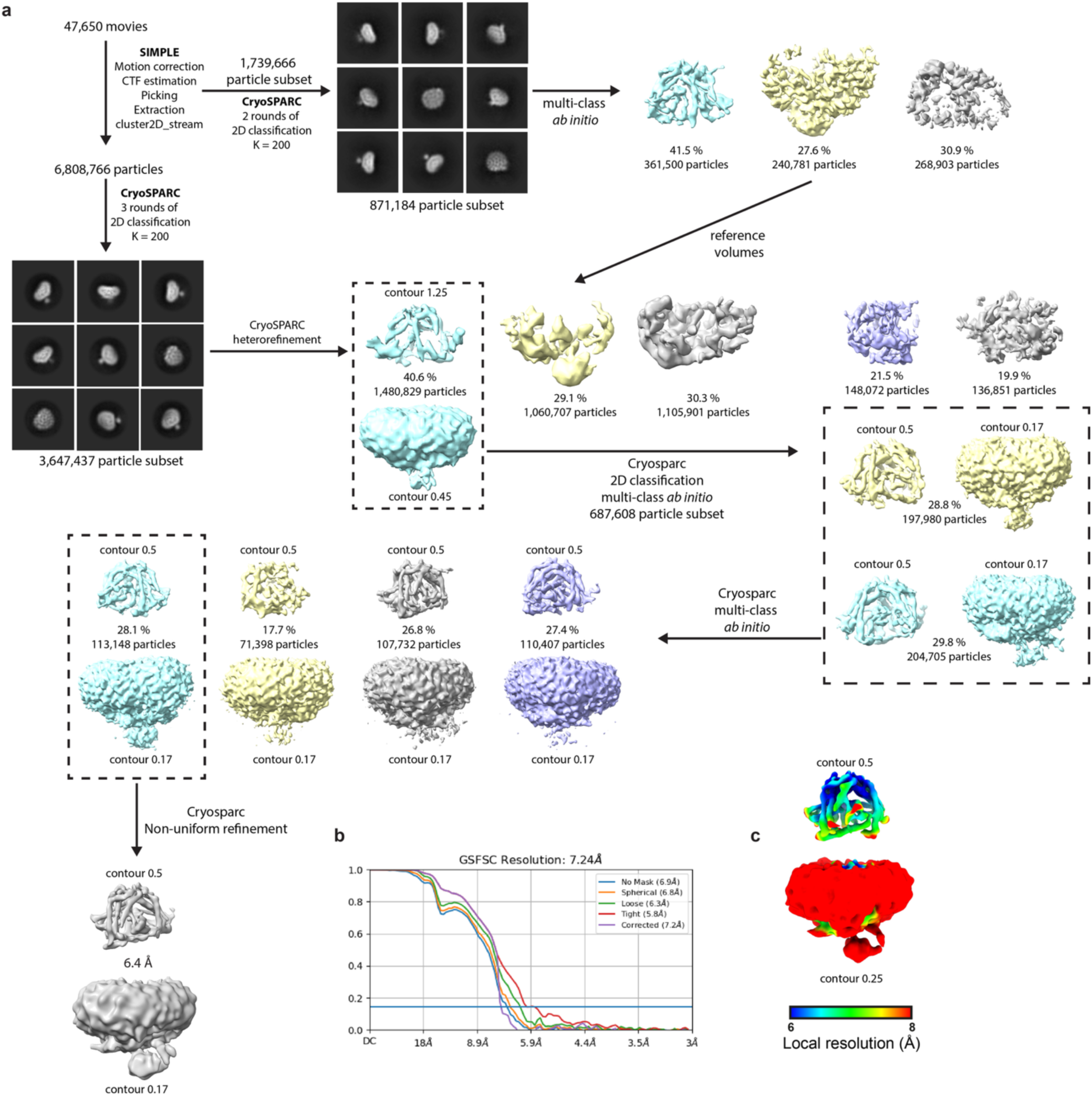
Cryo-EM workflow for an *E. coli* TatA_ALFA_BC-nanobody complex. **a**, Image processing workflow. **b**, Gold-standard Fourier Shell Correlation (FSC) curves used for global resolution estimation. **c**, Local resolution estimation of the volume

**Fig. S15.**
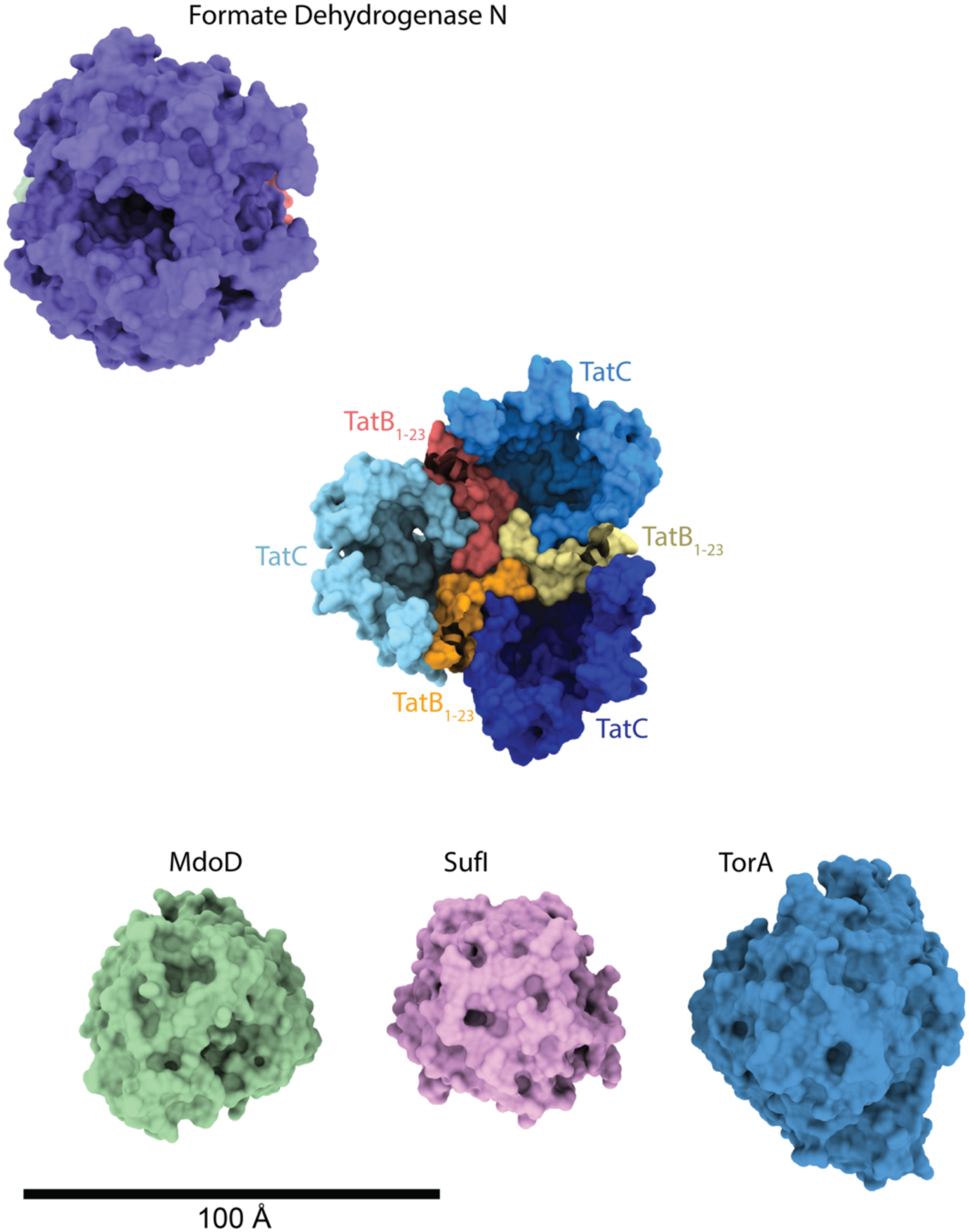
Sizes of the folded domains of a selection of Tat substrates relative to the size of the TatBC complex. A surface representation of TatBC viewed from the cytoplasm is shown in the centre of the figure (coloured as in main text figures) with the amphipathic helices of TatB removed. Surface representations of four substrates are shown below – with each substrate oriented to present the minimal cross-section for transport. None of these substrates could be accommodated within the TatBC complex without structural rearrangement that would essentially equate to complete disassembly of the TatBC complex.

---

**Table S1.**
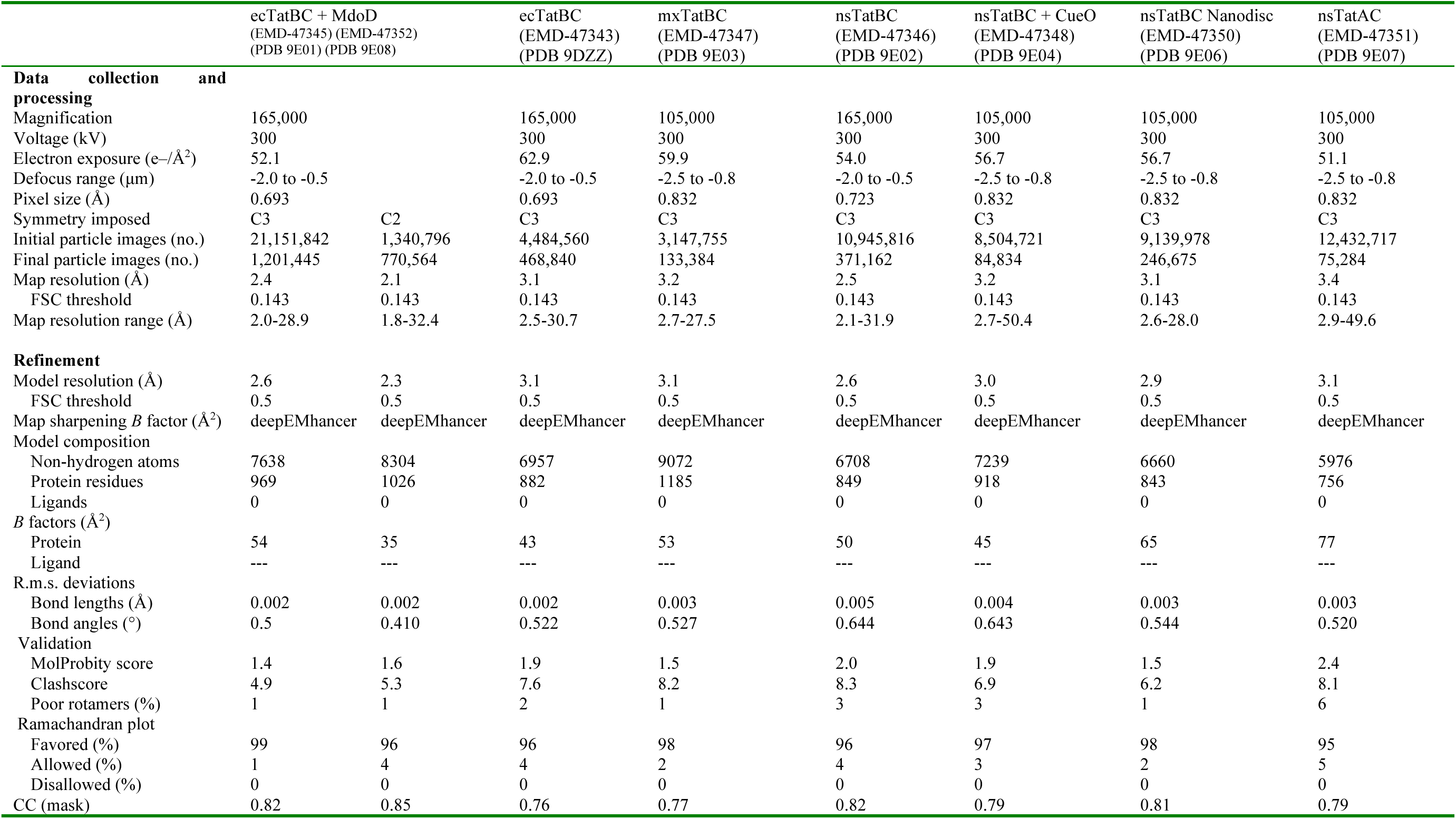
Cryo-EM data collection, refinement, and validation statistics.

|  | ecTatBC + MdoD<br>(EMD-47345) (EMD-47352)<br>(PDB 9E01) (PDB 9E08) |  | ecTatBC<br>(EMD-47343)<br>(PDB 9DZZ) | mxTatBC<br>(EMD-47347)<br>(PDB 9E03) | nsTatBC<br>(EMD-47346)<br>(PDB 9E02) | nsTatBC + CueO<br>(EMD-47348)<br>(PDB 9E04) | nsTatBC Nanodisc<br>(EMD-47350)<br>(PDB 9E06) | nsTatAC<br>(EMD-47351)<br>(PDB 9E07) |
| --- | --- | --- | --- | --- | --- | --- | --- | --- |
| **Data collection and processing** |  |  |  |  |  |  |  |  |
| Magnification | 165,000 |  | 165,000 | 105,000 | 165,000 | 105,000 | 105,000 | 105,000 |
| Voltage (kV) | 300 |  | 300 | 300 | 300 | 300 | 300 | 300 |
| Electron exposure (e-/Å²) | 52.1 |  | 62.9 | 59.9 | 54.0 | 56.7 | 56.7 | 51.1 |
| Defocus range (µm) | -2.0 to -0.5 |  | -2.0 to -0.5 | -2.5 to -0.8 | -2.0 to -0.5 | -2.5 to -0.8 | -2.5 to -0.8 | -2.5 to -0.8 |
| Pixel size (Å) | 0.693 |  | 0.693 | 0.832 | 0.723 | 0.832 | 0.832 | 0.832 |
| Symmetry imposed | C3 | C2 | C3 | C3 | C3 | C3 | C3 | C3 |
| Initial particle images (no.) | 21,151,842 | 1,340,796 | 4,484,560 | 3,147,755 | 10,945,816 | 8,504,721 | 9,139,978 | 12,432,717 |
| Final particle images (no.) | 1,201,445 | 770,564 | 468,840 | 133,384 | 371,162 | 84,834 | 246,675 | 75,284 |
| Map resolution (Å) | 2.4 | 2.1 | 3.1 | 3.2 | 2.5 | 3.2 | 3.1 | 3.4 |
| FSC threshold | 0.143 | 0.143 | 0.143 | 0.143 | 0.143 | 0.143 | 0.143 | 0.143 |
| Map resolution range (Å) | 2.0-28.9 | 1.8-32.4 | 2.5-30.7 | 2.7-27.5 | 2.1-31.9 | 2.7-50.4 | 2.6-28.0 | 2.9-49.6 |
| **Refinement** |  |  |  |  |  |  |  |  |
| Model resolution (Å) | 2.6 | 2.3 | 3.1 | 3.1 | 2.6 | 3.0 | 2.9 | 3.1 |
| FSC threshold | 0.5 | 0.5 | 0.5 | 0.5 | 0.5 | 0.5 | 0.5 | 0.5 |
| Map sharpening *B* factor (Å²) | deepEMhancer | deepEMhancer | deepEMhancer | deepEMhancer | deepEMhancer | deepEMhancer | deepEMhancer | deepEMhancer |
| Model composition |  |  |  |  |  |  |  |  |
| Non-hydrogen atoms | 7638 | 8304 | 6957 | 9072 | 6708 | 7239 | 6660 | 5976 |
| Protein residues | 969 | 1026 | 882 | 1185 | 849 | 918 | 843 | 756 |
| Ligands | 0 | 0 | 0 | 0 | 0 | 0 | 0 | 0 |
| *B* factors (Å²) |  |  |  |  |  |  |  |  |
| Protein | 54 | 35 | 43 | 53 | 50 | 45 | 65 | 77 |
| Ligand | --- | --- | --- | --- | --- | --- | --- | --- |
| R.m.s. deviations |  |  |  |  |  |  |  |  |
| Bond lengths (Å) | 0.002 | 0.002 | 0.002 | 0.003 | 0.005 | 0.004 | 0.003 | 0.003 |
| Bond angles (°) | 0.5 | 0.410 | 0.522 | 0.527 | 0.644 | 0.643 | 0.544 | 0.520 |
| Validation |  |  |  |  |  |  |  |  |
| MolProbity score | 1.4 | 1.6 | 1.9 | 1.5 | 2.0 | 1.9 | 1.5 | 2.4 |
| Clashscore | 4.9 | 5.3 | 7.6 | 8.2 | 8.3 | 6.9 | 6.2 | 8.1 |
| Poor rotamers (%) | 1 | 1 | 2 | 1 | 3 | 3 | 1 | 6 |
| Ramachandran plot |  |  |  |  |  |  |  |  |
| Favored (%) | 99 | 96 | 96 | 98 | 96 | 97 | 98 | 95 |
| Allowed (%) | 1 | 4 | 4 | 2 | 4 | 3 | 2 | 5 |
| Disallowed (%) | 0 | 0 | 0 | 0 | 0 | 0 | 0 | 0 |
| CC (mask) | 0.82 | 0.85 | 0.76 | 0.77 | 0.82 | 0.79 | 0.81 | 0.79 |

**Table S2.** Data collection, phasing and refinement statistics for *M. xanthus* periplasmic domain.

|  | SeMet derivatized crystal#<br>9E0S |
| --- | --- |
| **Data collection** |  |
| Space group | P422 |
| Cell dimensions |  |
| *a*, *b*, *c* (Å) | 110.83, 110.83, 102.13 |
| $\alpha$, $\beta$, $\gamma$ (°) | 90,90,90 |
|  | *Peak* |
| Wavelength | 0.97926 |
| Resolution (Å) | 62.1-1.98 (2.03-1.98) |
| *R*<sub>pim</sub> | 0.054 (0.521) |
| *I* / $\sigma I$ | 11.7 (1.7) |
| Completeness (%) | 100 (100) |
| Redundancy | 25.2 (25.6) |
| **Refinement** |  |
| Resolution (Å) | 62.1-1.98 (2.03-1.98) |
| No. reflections | 44,477 (3,065) |
| *R*<sub>work</sub> / *R*<sub>free</sub> | 0.19 / 0.22 (0.26 / 0.30) |
| No. atoms | 4365 |
| Protein residues | 529 |
| Ligand/ion | 1 |
| Water | 321 |
| *B*-factors |  |
| Protein | 31 |
| Ligand/ion | 21 |
| Water | 33 |
| R.m.s deviations |  |
| Bond lengths (Å) | 0.007 |
| Bond angles (°) | 0.8 |
| Validation |  |
| MolProbity score | 1.4 |
| Clashscore | 3.7 |
| Poor rotamers (%) | 2 |
| Ramachandran plot |  |
| Favored (%) | 99 |
| Allowed (%) | 1 |
| Disallowed (%) | 0 |

#A single crystal was used for data collection. \*Values in parentheses are for highest-resolution shell.

|  | Genotype |
| --- | --- |
| L56 (DE3) | *REFERENCE* |
| BL21 (DE3) $\Delta tatABCD$, $\Delta tatE$ | B F<sup>-</sup> *ompT gal dcm lon hsdS<sub>B</sub>(r<sub>B</sub><sup>-</sup> m<sub>B</sub><sup>-</sup>)* $\lambda$(DE3 [*lacI lacUV5-T7p07 ind1 sam7 nin5*]) [*malB*<sup>+</sup>]<sub>K-12</sub>($\lambda^S$), $\Delta tatABCD$ |
| MC4100-A | F<sup>-</sup>, [*araD139/B/r*, $\Delta$(*argF-lac*)169, $\lambda^-$, *e14-*, *flhD5301*, $\Delta$(*fruK-yeiR*)725(*fruA25*), *relA1*, *rpsL150*(strR), *rbsR22*, $\Delta$(*fimB-fimE*)632(*::IS1*), *deoC1*, arabinose resistant |
| MC4100-A $\Delta tatABCD$, $\Delta tatE$ | MC4100-A, $\Delta tatABCD$, $\Delta tatE$ |
| MC4100-A $\Delta mdoD$ | MC4100-A, $\Delta mdoD$ |
| MC4100-A tatC:strep | MC4100-A, *tatC*<sub>strep</sub> |
| MC4100-A tatC:strep L99S | MC4100-A, *tatC*:strep, *tatC*<sub>L99S</sub> |
| MC4100-A tatC:strep E103A | MC4100-A, *tatC*<sub>E103A</sub> |
| B834 (DE3) | F-*ompT hsdSB*(rB- mB-) *gal dcm met* (DE3) |

| Plasmid | Description |
| --- | --- |
| Mx-pTatCperi-His | pET15b based vector carrying *Myxococcus xanthus* TatC periplasmic domain sequence with C-terminal TEV protease site and hexahistidine tag. |
| pFAT75ΔA | pQE60 based vector carrying *Escherichia coli* *tatBC* encoding a C-terminal histidine tag. |
| pUNITAT2 | pQE60 based vector carrying *Escherichia coli* *tatABC* encoding a C-terminal histidine tag. |
| pREP4 | Vector carrying lacI gene. |
| Mx-pTatC-GFP-His | pWALDO based vector carrying *Myxococcus xanthus* TatC containing sequence encoding a C-terminal GFP and hexa-histidine tag. |
| Mx-pTatBstrep | pCDFDuet-1 based vector carrying *Myxococcus xanthus* TatB containing sequence encoding a C-terminal strep tag. |
| Ns-pTatC-GFP-His | pWALDO based vector carrying *Nitratifractor salsuginis* TatC containing sequence encoding a C-terminal GFP and hexa-histidine tag. |
| Ns-pTatBstrep | pCDFDuet-1 based vector carrying *Nitratifractor salsuginis* TatB containing a C-terminalstrep tag. |
| Ns-pTatAstrep | pCDFDuet-1 based vector carrying *Nitratifractor salsuginis* TatA containing sequence encoding a C-terminal strep tag. |
| Ns-pTatCstrep | pRSFDuet-1 based vector carrying *Nitratifractor salsuginis* TatC containing sequence encoding a C-terminal strep tag. |
| Ns-pTatA_His | pACT7 based vector carrying *Nitratifractor salsuginis* TatA containing sequence encoding a C-terminal hexa-histidine tag. |
| pUNITAT2strep TatA_His | pQE60 based vector carrying *Escherichia coli* *tatABC* containing sequence encoding a C-terminal strep tag and sequence to introduce a hexa-histidine tag between tatA codons 26 and 27. |
| pUNITAT2strep TatA_His_ALFA | pQE60 based vector carrying *Escherichia coli* *tatABC* containing sequence encoding a C-terminal strep tag and sequence to introduce an ALFA tag followed by a hexa-histidine tag between tatA codons 23 and 24. |
| Ec-pTatA | pQE80 based vector carrying *Escherichia coli* *tatA*. |
| Ec-pTatB | pQE80 based vector carrying *Escherichia coli* *tatB*. |
| Ns-pTatCstrep-TatB1flag | PRSFduet-1 based vector carrying *Nitratifractor salsuginis* TatC containing sequence encoding a C-terminal strep tag inserted into mcs1 and *N. salsuginis* TatB containing C-terminal flag tag in mcs2.. |
| Ns-pTatCstrep-L99S-TatB1flag | PRSFduet-1 based vector carrying *Nitratifractor salsuginis* TatC containing sequence encoding a C-terminal strep tag inserted into mcs1 and *N. salsuginis* TatB containing C-terminal flag tag in mcs2. |
| Ns-pTatCstrep-E103A-TatB1flag | PRSFduet-1 based vector carrying *Nitratifractor salsuginis* TatC containing sequence encoding a C-terminal strep tag inserted into mcs1 and *N. salsuginis* TatB containing C-terminal flag tag in mcs2. |
| pET28a_(His)6-SUMO-MdoD | pET28a based vector carrying *Escherichia coli mdoD* containing an N-terminal hexa-histidine-SUMO fusion.= |
| pET28a_(His)6-SUMO-CueO | pET28a based vector carrying *Escherichia coli cueO* containing an N-terminal hexa-histidine-SUMO fusion. |
| pQE80_MdoD-(His)6 | pBAD33 based vector carrying *Escherichia coli mdoD* containing a C-terminal hexa-histidine tag. |
| pBAD33 | pBAD33 CITE |
| pBAD33_MdoDΔ1-32-myc | pQE80 based vector carrying *Escherichia coli mdoD* with the sequence encoding the Tat export signal deleted (corresponding to residues 1-32). |
| pQE80_CueO-(His)6 | pQE80 based vector carrying *Escherichia coli cueO* with sequence encoding a C-terminal hexa-histidine tag. |
| pQE80_MdoD_myc | pQE80 based vector carrying *Escherichia coli mdoD* with sequence encoding a C-terminal myc tag. |
| pQE80_MdoD_myc Δ28-43 | pQE80 based vector carrying *Escherichia coli mdoD* with sequence encoding a C-terminal myc tag. Sequence encoding residues 28-43 is deleted. |
| pQE80_MdoD_myc Δ28-46 | pQE80 based vector carrying *Escherichia coli mdoD* with sequence encoding a C-terminal myc tag. Sequence encoding residues 28-46 is deleted. |
| pWRG730 | Temperature sensitive plasmid encoding λ-red phage genes (*gam*, *bet*, *exo*). |
| pWRG717 | Plasmid carrying kanamycin resistance gene (*aph*) that contains Sce-I restriction site. Used as template to produce a PCR product to replace genes for scarless mutagenesis. |

